# Anti-viral defense by an ADP-ribosyltransferase that targets mRNA to block translation

**DOI:** 10.1101/2024.02.24.581662

**Authors:** Christopher N. Vassallo, Christopher R. Doering, Michael T. Laub

## Abstract

Host-pathogen conflicts are crucibles of molecular innovation. Selection for immunity to pathogens has driven the evolution of sophisticated immunity mechanisms throughout biology, including in bacteria that must evade their viral predators known as bacteriophages. Here, we characterize a widely distributed anti-phage defense system, CmdTAC, that provides robust defense against infection by the T-even family of phages. Our results support a model in which CmdC detects infection by sensing viral capsid proteins, ultimately leading to the activation of a toxic ADP-ribosyltransferase effector protein, CmdT. We show that newly synthesized capsid protein triggers dissociation of the chaperone CmdC from the CmdTAC complex, leading to destabilization and degradation of the antitoxin CmdA, with consequent liberation of the CmdT ADP-ribosyltransferase. Strikingly, CmdT does not target a protein, DNA, or structured RNA, the known targets of other ADP-ribosyltransferases. Instead, CmdT modifies the N6 position of adenine in GA dinucleotides within single-stranded RNAs leading to arrest of mRNA translation and the inhibition of viral replication. Our work reveals a new mechanism of anti-viral defense and a previously unknown but broadly distributed class of ADP-ribosyltransferases that target mRNA.

## Introduction

ADP-ribosyltransferases are important enzymes found throughout biology^1–3^. These enzymes transfer the ADP-ribose moiety of NAD^+^ onto other biomolecules, usually modifying an amino acid on a target protein. In some cases, these enzymes are mono-ADP ribosyltransferases (mARTs), adding a single ADP-ribose group to a protein while in other cases they are poly-ADP-ribosyl polymerases (PARPs) that can add ADP-ribose groups to the 2’ OH on the adenine of an existing ADP-ribose modification, leading to polymers of ADP-ribose. ADP-ribosyltransferase activity was first documented in liver cell extracts^4^, with subsequent work demonstrating that a toxic protein secreted by *Bordetella pertussis* ADP-ribosylates human EF2 to block translation in host cells^5^. In the ensuing decades, extensive work revealed ADP-ribosylation as one of the most common post-translational modifications in biology, used to regulate a wide variety of proteins involved in cellular signaling, chromatin and transcription, DNA repair, and more. At least 6 of the human PARPs are interferon-induced and thus hypothesized to function in anti-viral defense^6,7^, but their precise roles and targets remain poorly defined.

Although studied for decades as protein modifiers, recent work identified ADP-ribosyltransferases that target nucleic acids. DarTG toxin-antitoxin systems include a toxin, DarT, that can modify either thymidine or guanosine, depending on the DarT homolog, in single-stranded DNA^8,9^. Some DarTG systems have potent anti-phage defense activity, with bacteriophage infection leading to release of DarT from its cognate DarG antitoxin, modification of phage DNA, and a block in phage DNA replication^10^. In *Pseudomonas aeruginosa*, the type VI secretion system effector RhsP2 kills cells by ADP-ribosylating the 2’-OH of some double-stranded RNAs, though the precise RNA target leading to cell death is unclear^11^. In *Photorhabdus laumondii* a type VI secretion system effector modifies 23S rRNA to block translation^12^.

Here, we discover an ADP-ribosyltransferase called CmdT that, like DarT, functions in bacterial anti-phage defense, but does so by specifically modifying mRNAs to block phage translation and the production of mature virions. CmdT is part of a tri-partite toxin-antitoxin-chaperone (TAC) system. Toxin-antitoxin (TA) systems are extremely prevalent in bacterial genomes and play a major role in anti-phage defense^13^. TA systems feature a protein toxin that is restrained from killing a cell or blocking cell growth by a cognate antitoxin^14^. For anti-phage TA systems, phage infection must somehow liberate the toxin, but the mechanisms responsible remain incompletely understood^15,16^. TAC systems are common variants that feature a chaperone related to the conserved protein export chaperone, SecB. The chaperones of TAC systems promote proteolytic stability of their cognate antitoxin and, consequently, neutralization of the toxin^17–20^. How the toxins of TAC systems are liberated is not known.

For CmdTAC from the *Escherichia coli* strain ECOR22 (ref^21^), we find that, during infection by T4 phage, the major capsid protein outcompetes the antitoxin, CmdA, for binding to the chaperone, CmdC. Consequently, CmdA is degraded by the ClpP protease, leading to activation of the CmdT ADP-ribosyltransferase. Strikingly, CmdT does not target proteins, DNA, or structured RNAs, the targets of other known ADP-ribosyltransferases. Instead, we find that CmdT selectively modifies mRNA to block translation and abort the phage infection. Biochemical analyses demonstrate that CmdT primarily modifies the N6 position of adenine in GA dinucleotides found in mRNAs. In sum, our work reveals both a novel mechanism of anti-phage defense and a previously unknown molecular target of an ADP-ribosyltransferase. As some of the interferon-inducible human PARPs are reported to also modify RNA *in vitro*^7^, our work suggests that ADP-ribosylation of RNA may be a common, previously unappreciated facet of innate immunity throughout biology.

## Results

### CmdTAC protects against *Tevenirinae* phages by an abortive infection mechanism

The *cmdTAC* system (Fig. 1A) was discovered in the genome of the wild *E. coli* isolate ECOR22 though its homologs are found in diverse bacteria, including both Gram-negative and Gram-positive species (Fig. S1A-B). In *E. coli*, *cmdTAC* provides robust defense against a major family of phages, the *Tevenvirinae*, which includes T4. Consistent with our prior work^21^, deleting *cmdTAC* from ECOR22 improved the efficiency of plaquing (EOP) of T4 by at least 10^3^-fold. Transformation of ECOR22 with a plasmid containing *cmdTAC* expressed from its native promoter fully restored defense (Fig. 1B). We previously found that plasmid-based *cmdTAC* in *E. coli* MG1655 provided defense against the T-even phages T2, T4, and T6 (ref^21^). Here, we challenged *E. coli* MG1655 harboring *cmdTAC* with all 12 *Myoviridae* subfamily *Tevenirinae* phage from the BASEL collection^22^. CmdTAC strongly decreased the EOP of most *Tevenirinae* phages with only two exceptions, Bas35 and Bas38 (Fig. 1C, Fig. S1C). No protection was observed against phages from other taxa. To directly assess the impact of CmdTAC on viral progeny production, we measured the burst size of T4 after a single round of infection. The presence of CmdTAC reduced the number of T4 progeny produced from ∼110 phages to almost 0, indicating a lack of new viral particle production (Fig. 1D).

**Figure 1.**
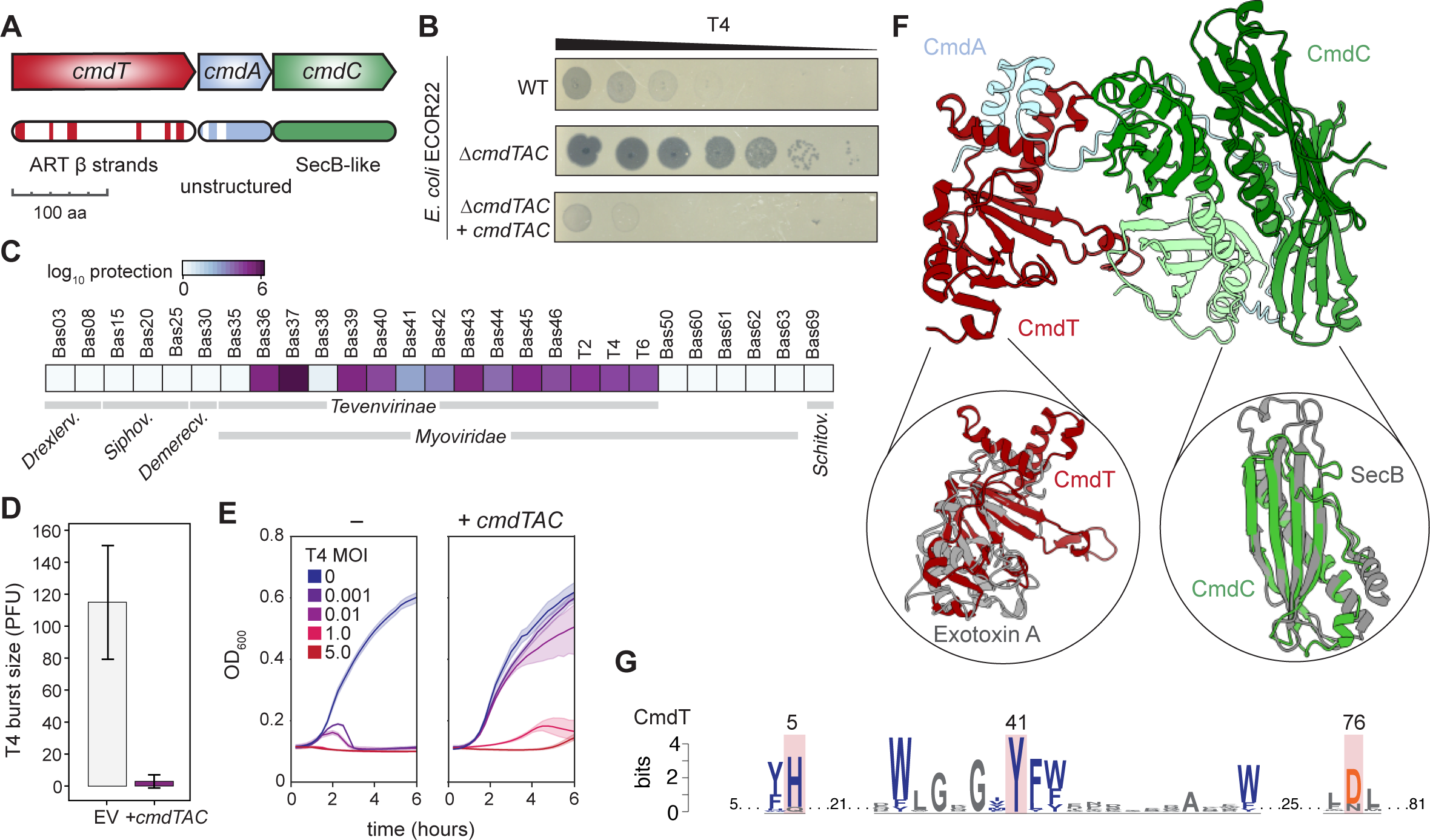
The anti-phage defense system CmdTAC protects *E. coli* against infection by *Tevenviridae*. (A) Schematic of the *cmdTAC* operon indicating predicted structural features. ART = ADP-ribosyltransferase. (B) Plaquing of 10-fold serially diluted T4 phage on lawns of wild-type (WT) *E. coli* strain ECOR22 and the Δ*cmdTAC* mutant without or with a complementing plasmid. (C) Efficiency of plaquing of the phages indicated on *E. coli* K12 harboring *cmdTAC* relative to an empty vector control. *Drexlerv., Siphov., Demerecv., and Schitov.,* are abbreviations for *Drexlerviridae, Siphoviridae, Demerecviridae,* and *Schitoviridae* respectively. T2, T4, and T6 data taken from ref^21^. (D) T4 burst size in *E. coli* K12 harboring *cmdTAC* or an empty vector. Error bars represent 95% confidence interval for five biological replicates. (E) Growth of *E. coli* K12 harboring *cmdTAC* or an empty vector and infected with T4 at the indicated MOIs. (F) AlphaFold2-predicted model of CmdTAC complex with insets showing structural alignments of CmdT to the ADP-ribosyltransferase Exotoxin A (PBD 1AER, RMSD 6.93) and CmdC to the SecB chaperone (PBD 1OZB, RMSD 4.13). (G) Sequence logo showing conservation of key catalytic residues in CmdT homologs. Shaded residues indicate active site residues in known ADP-ribosyltransferases. Blue color indicates conserved, aromatic residues. Amino acid numbers for CmdT^ECOR22^ listed across the top.

To determine how CmdTAC provides defense, we examined cell growth of *E. coli* MG1655 containing *cmdTAC* on a plasmid under its native promoter following infection by T4 at different multiplicities of infections (MOI) (Fig. 1E). Direct defense systems provide protection at almost any MOI. In contrast, abortive infection (Abi) systems allow growth in the presence of phage only at MOIs less than 1.0. This is because Abi systems either inhibit growth of infected cells or block a phage from replicating after the phage triggers host cell death and thus function only to prevent the spread of an infection through the population^23^. We observed protection against T4 infection only at low (< 1) MOIs, indicating that CmdTAC functions through an Abi mechanism^21^.

### CmdA promotes the folding and neutralization of CmdT but is proteolytically unstable

To identify the potential functions of each component in the CmdTAC system, we performed remote homology searches using HHpred^24^ on each protein and used AlphaFold2 (ref^25^) to predict the structures of the components separately and in complex (Fig. 1F and Fig. S2). These analyses identified ADP-ribosyltransferase-like β-strands in CmdT and predicted a structure with similarity to the catalytic domain of *P. aeruginosa* exotoxin A, an ADP-ribosyltransferase of the DTX family that transfers the ADP-ribose moiety of nicotinamide adenine dinucleotide (NAD^+^) onto the eukaryotic translation factor eEF2 (ref^26^). Building a sequence logo of CmdT homologs also revealed several conserved residues including the HYE/D catalytic triad of the DTX family of ADP-ribosyltransferases^27^ (Fig. 1G). Substituting alanine for the inferred catalytic residue of CmdT, Y41, (a variant referred to here as CmdT*), abolished protection against T4 (Fig. S1D). HHpred did not yield any predictions for CmdA, but AlphaFold2 predicted a pair of N-terminal α-helices followed by a long and unstructured C-terminal domain. CmdC is homologous to SecB and has the same predicted tetrameric organization as *E. coli* SecB and the SecB-like chaperones commonly found in toxin-antitoxin-chaperone (TAC) systems^18,19^. AlphaFold2 predicted with high confidence (Fig. S2) a hetero-hexameric complex in which the N-terminal α-helices of CmdA interface with α-helices in CmdT while the unstructured C-terminus of CmdA stretches from the putative catalytic pocket of CmdT to a binding interface with tetrameric CmdC (Fig. 1F).

To experimentally test the predicted functions of the CmdTAC components, we cloned each gene individually under an inducible promoter in *E. coli* MG1655. Inducing any individual component did not strongly affect plating viability (Fig. 2A). However, co-expressing *cmdT* and *cmdA* led to a substantial loss in viability. This phenotype was abolished by the Y41A substitution in the highly conserved tyrosine in the putative catalytic domain of CmdT, indicating that CmdT is likely a toxic ADP-ribosyltransferase (Fig. 2A). We considered two hypotheses to explain the dependency of CmdT toxicity on the co-expression of CmdA: (1) CmdT and CmdA together form an active effector or (2) CmdA is required for the production or stabilization of active CmdT. To distinguish between these hypotheses, we first used immunoblotting to measure the levels of FLAG-tagged CmdT produced alone or together with CmdA from an inducible promoter. Strikingly, when produced alone, CmdT was not detectable on a native gel lacking SDS, whereas CmdT co-produced with CmdA was present and increased in abundance over time (Fig. 2B, Fig. S3A). This result indicated that CmdT was either not present, or aggregated and could not enter the native gel. Using the same samples, but with denaturing SDS-PAGE, we found CmdT produced alone was detectable, but with fluctuating levels over time. In contrast, CmdT co-produced with CmdA readily accumulated. These results suggest that CmdT produced alone aggregated and is non-functional, but can stably fold when co-produced with CmdA.

**Figure 2.**
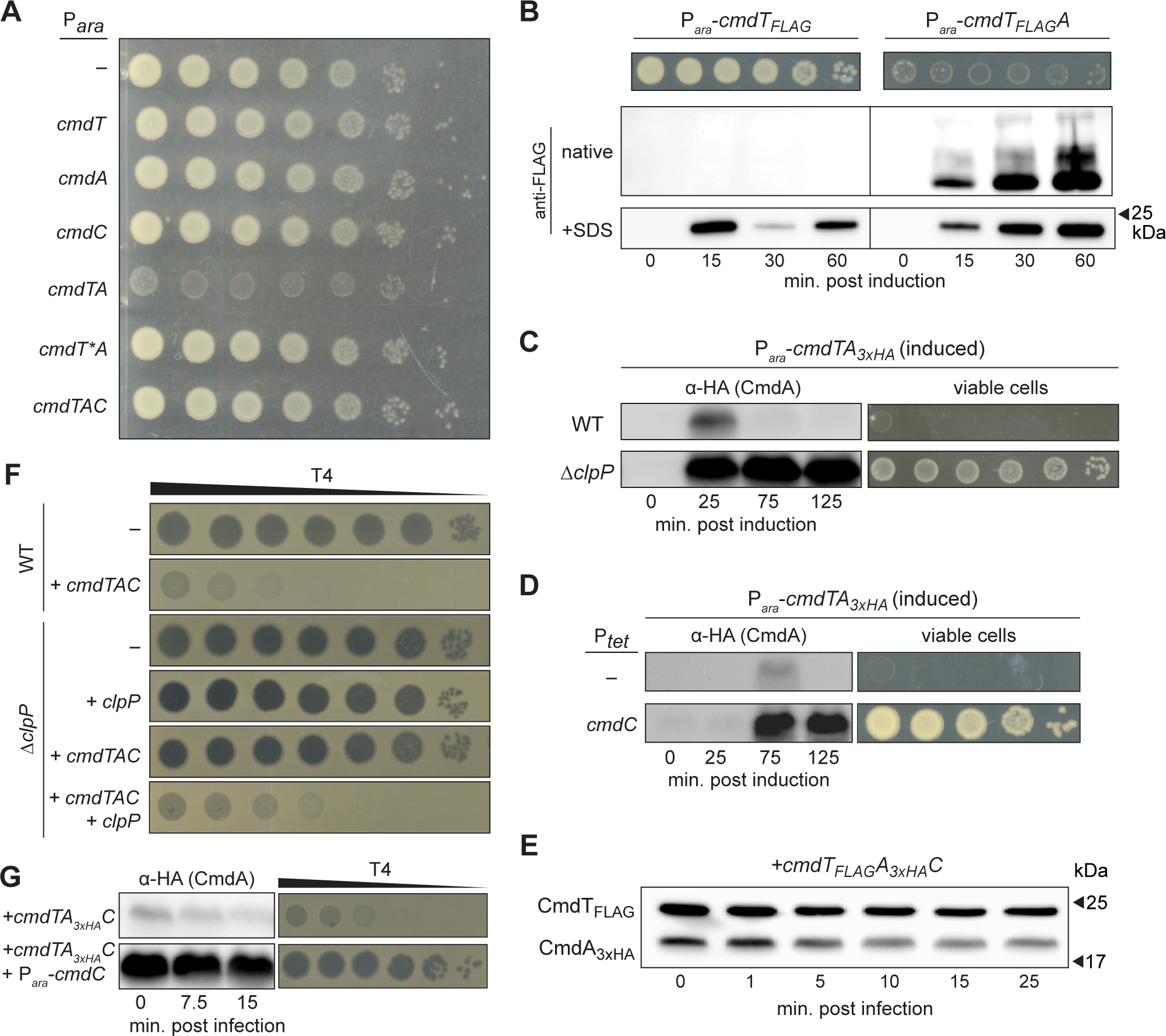
CmdTAC is a toxin-antitoxin-chaperone system. (A) Plating viability (10-fold serial dilutions) of strains inducibly expressing the components listed. *cmdT\** harbors a point mutation predicted to ablate ADP-ribosyltransferase activity. (B) Cells harboring *cmdT* or *cmdTA*, each with a FLAG tag on the C-terminus of CmdT, were induced with assessment of plating viability (top) and CmdT levels via immunoblotting of native gels (middle) or standard SDS-PAGE (bottom). Coomassie straining used as an approximate loading control is in Fig. S3A. (C) Wild-type or Δ*clpP* cells expressing *cmdTA* with a 3xHA tag on the N-terminus of CmdA, with assessment of CmdA levels via immunoblotting (left) and plating viability (right). (D) Cells expressing *cmdTA* from an inducible promoter with CmdA engineered to have an N-terminal 3xHA tag. Cells also express *cmdC* from a tetracycline-inducible promoter. CmdA levels were measured by immunblot (left), with cell viability assessed by 10-fold serial dilutions (right). (E) Immunoblot, using anti-FLAG and anti-HA antibodies, of cells harboring *cmdTAC* expressed from its native promoter with CmdT and CmdA engineered to have a FLAG or 3xHA tag, respectively, on their N-termini. Samples taken from cells infected with T4 for the indicated times. Coomassie staining used as an approximate loading control is in Fig. S3B. (F) Plaquing of 10-fold serially diluted T4 phage on lawns of the strains indicated. (G) Cells harboring *cmdTAC* under native promoter control with CmdA engineered to have an N-terminal 3xHA tag and expressing an additional copy of *cmdC* (bottom) or not (top) with assessment of CmdA levels by immunoblot (left) and anti-phage defense by 10-fold serially diluted T4 plaquing (right).

Inducing *cmdTA* is toxic in the absence of CmdC. We predicted that this toxicity occurs because CmdA is proteolytically degraded without its CmdC chaperone. To test this prediction, we repeated the *cmdTA* co-expression experiment, but with CmdA harboring a 3xHA epitope, and found that CmdA transiently accumulated and then was almost undetectable after 75 minutes of induction (Fig. 2C). Taken all together, our results indicate that when co-expressing *cmdTA*, CmdA is initially needed for the folding or stabilization of CmdT, but that CmdA is subsequently cleared from cells, leaving behind stable, toxic CmdT.

Like the antitoxins of other known TAC systems, the C-terminus of CmdA ends in ‘LAA’, a known ClpXP recognition sequence^28^. To determine if CmdA is degraded by ClpP, we introduced an inducible version of *cmdTA_3xHA_* into a *ΔclpP* strain and again monitored CmdA levels by immunoblot. In comparison to the wild-type background where CmdA accumulates and then is undetectable, CmdA accumulated to consistently high levels in the absence of ClpP (Fig. 2C). This accumulated CmdA was sufficient to neutralize the toxicity of CmdT as reflected by the substantially improved plating viability of *ΔclpP* cells co-expressing *cmdTA* compared to wild-type cells co-expressing *cmdTA*.

We also found that co-expressing *cmdC*, either in the same operon as *cmdTA* (Fig. 2A) or separately (Fig. 2D) was sufficient to rescue the lethality of co-expressing *cmdTA*, and immunoblotting confirmed that CmdC production promoted the accumulation of CmdA (Fig. 2D). Collectively, our findings indicate that CmdTAC is a toxin-antitoxin-chaperone system with the following features. CmdT is a toxin that requires CmdA to fold and stably accumulate. CmdA also serves as an antitoxin to neutralize CmdT in the presence of the SecB-like chaperone CmdC. In the absence of CmdC, CmdA is degraded by the ClpP protease, leading to CmdT toxicity.

### CmdA degradation is necessary for CmdTAC-based phage defense

In the absence of phage infection, CmdC and CmdA neutralize CmdT. Upon phage infection, CmdT must somehow be released to abort the infection and prevent the production of new viruses. To understand the dynamics of the CmdTAC system, we used T4 to infect cells harboring a variant of *cmdTAC* that produces 3xHA-tagged CmdA and FLAG-tagged CmdT; we then monitored the presence of each component through immunoblotting. CmdT was present throughout the course of phage infection, whereas CmdA levels steadily decreased (Fig. 2E, Fig. S3B).

The decrease in CmdA during phage infection likely reflects its degradation by ClpXP and suggests that the ClpP protease is essential for CmdT activation and phage defense. To test this hypothesis, we transformed a plasmid containing the *cmdTAC* system under control of its native promoter, or an empty vector, into a *ΔclpP* strain and then challenged cells with T4 phage. We found that T4 robustly infected *ΔclpP* cells harboring either *cmdTAC* or an empty vector, in stark contrast to WT cells harboring *cmdTAC* that exhibited strong defense against T4 (Fig. 2F). The defect in defense manifested by the *ΔclpP* strain was largely reversed by introducing a copy of *clpP* in trans.

As a further test that CmdA degradation is necessary for CmdT-based defense, we co-transformed cells with a plasmid containing *cmdTAC* (with *cmdA* containing a 3xHA epitope tag) and a plasmid harboring a second copy of *cmdC* under control of an arabinose-inducible promoter. Overproducing CmdC led to the accumulation of CmdA and a loss of defense against T4 (Fig. 2G). Taken together, our results indicate that CmdTAC likely forms a stable complex in the absence of phage, with infection then triggering the ClpP-dependent degradation of CmdA, release of CmdT, and an aborted infection.

### T4 can escape CmdTAC by blocking ClpP proteolysis of CmdA

How does phage infection trigger CmdA degradation and consequent liberation of CmdT? To answer this question, we selected escape mutants of T4 that can evade CmdTAC-based defense. Whole-genome sequencing of six independent isolates revealed that each escape clone harbored a mutation in the uncharacterized gene *alt.-3* that was either: (i) a single nucleotide deletion near the 3’ end of the coding region leading to a frameshift or (ii) a mutation in the normal stop codon (Fig. 3A). In each case, the mutation extended the Alt.-3 coding sequence to one of two alternative stop codons, leading to either an extended 110 or 117 amino acid long protein. To understand the effect of the mutations we selected the 117 amino acid long mutant for further experiments, referred to here as Alt.-3^†^. T4 producing Alt.-3^†^ had substantially improved plaquing on CmdTAC-containing cells, effectively abolishing the defense normally provided by CmdTAC (compare Fig. 3B to Fig. 2F).

**Figure 3.**
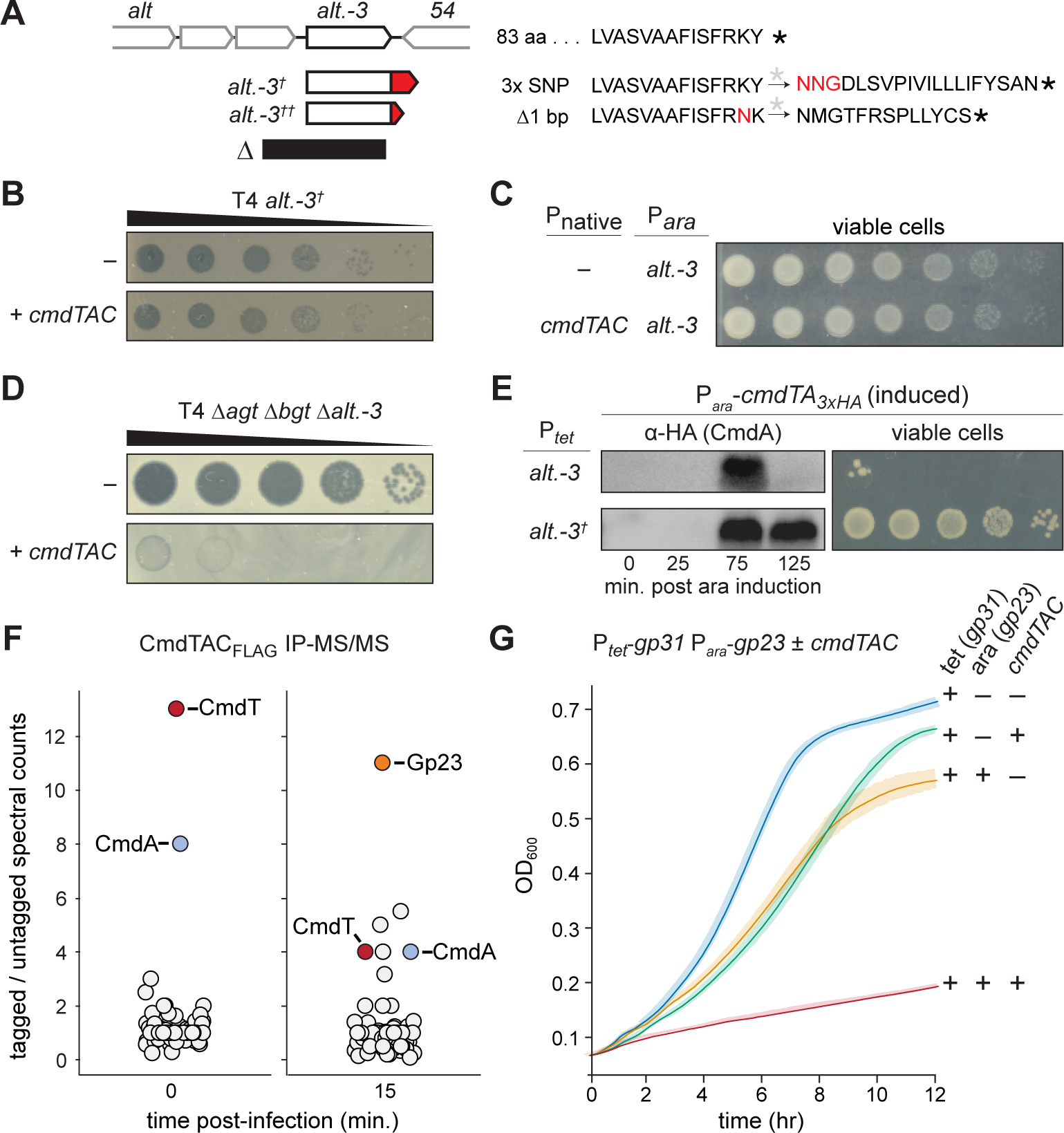
CmdTAC is activated by the T4 major capsid protein Gp23. (A) The *alt.-3*^†^ and *alt.-3*^††^ mutations isolated as *cmdTAC* escape mutants are summarized below a diagram of the *alt.-3* region of the T4 genome. The region deleted in T4 Δ*agt* Δ*bgt* Δ*alt.-3* is shown below. The C-termini of *alt.-3*, *alt.-3*^†^, and *alt.-3*^††^ are shown on the right. (B) Plaquing of 10-fold serially diluted T4 phage harboring the *alt.-3*^†^ mutation on lawns of *E. coli* expressing *cmdTAC* or carrying an empty vector. (C) Plating viability of strains harboring *cmdTAC* or an empty vector and expressing the wild-type *alt.- 3* gene. (D) Plaquing of 10-fold serially diluted T4 Δ*agt* Δ*bgt* Δ*alt.-3* phage on lawns of *E. coli* harboring *cmdTAC* or an empty vector. The bacterial genotype was Δ*mcrA* Δ*mcrBC*, as required for T4 Δ*agt* Δ*bgt*. (E) Cells expressing *cmdTA* from an inducible promoter with CmdA engineered to have an N-terminal 3xHA tag. Cells also express *alt.-3* or *alt.-3*^†^ from a tetracycline-inducible promoter. CmdA levels were measured by immunblot (left), with cell viability assessed by 10-fold serial dilutions (right). (F) Summary of IP-MS/MS analysis of proteins co-precipitating with N-terminally FLAG-tagged CmdC at 0 (left) or 15 (right) minutes after infecting cells harboring *cmdTAC* with T4 phage. Data points corresponding to CmdT, CmdA, and Gp23 are labeled. (G) Growth curves for uninfected *E. coli* cells producing CmdTAC from its native promoter, with induced production of Gp23 (T4 major capsid protein), and Gp31 (T4 chaperonin for Gp23), as indicated. Data are the average of 3 biological replicates.

To determine the basis of escape, we first considered whether Alt.-3 triggers the activation of the CmdTAC system, with the *alt.-3*^†^ mutations having prevented this activation. To test this possibility, we cloned *alt.-3* into a plasmid under inducible control and transformed it into cells harboring *cmdTAC*. However, producing Alt.-3 in the presence of CmdTAC caused no discernable loss in viability (Fig. 3C) indicating that Alt.-3 is not sufficient to activate CmdTAC. We also deleted *alt.-3* from the T4 genome but found that CmdTAC still provided robust defense against these phages (Fig. 3D). Thus, we ruled out Alt.-3 as an activator of CmdTAC.

Next, we hypothesized that the extended Alt.-3^†^ proteins somehow inhibited CmdTAC. To test this possibility, we asked whether producing Alt.-3 or Alt.-3^†^ could neutralize the toxicity of CmdTA. Inducing Alt.-3^†^, but not Alt.-3, resulted in a complete rescue of CmdTA toxicity (Fig. 3E, compare to Fig. 2G). This result could indicate either that Alt.-3^†^ directly inhibits CmdT toxicity or that it stabilizes CmdA, which then neutralizes CmdT. To differentiate between these possibilities, we co-produced Alt.- 3 or Alt.-3^†^ with CmdTA where CmdA harbors an HA epitope tag. Immunoblotting after Alt.-3^†^ induction revealed an increase and maintenance of CmdA levels comparable to that observed during CmdC induction (Fig. 3E, compare to Fig. 2D). Thus, Alt.-3^†^ inhibits CmdA degradation, promoting stabilization of the CmdTA complex and preventing activation of CmdTAC during phage infection. These results reinforce the conclusion that CmdT activation requires CmdA degradation and indicate that T4 phages can readily escape CmdTAC-mediated defense through mutations that inhibit this degradation.

### CmdTAC is activated by the T4 major capsid protein, Gp23

To identify the direct trigger for CmdTAC, we deleted *alt.-3* from the T4 genome (Fig. 3A) and attempted to isolate additional escape mutants but were unable to. Because CmdC is needed to protect CmdA from degradation (Fig. 2D), we suspected that during infection CmdC might preferentially bind a phage product over CmdA, leading to release and degradation of CmdA and consequent liberation of CmdT. In this way, CmdC could function as a sensor for the presence of a phage-encoded protein. To identify potential CmdC interacting partners, we FLAG-tagged CmdC in the context of *cmdTAC* under its native promoter and performed immunoprecipitation coupled to tandem mass spectrometry (IP-MS/MS) at 0 and 15 minutes post-T4 infection (Fig. 3F). As expected, before phage infection CmdC co-precipitated both CmdA and CmdT. However, at 15-minutes post infection, the major co-precipitating protein was Gp23, the major capsid protein of T4 (ref^29^).

To validate our IP-MS/MS result, we tested whether Gp23 is sufficient to activate CmdTAC and cause CmdT-dependent growth inhibition. We cloned *gp23* under an inducible promoter and transformed it into cells with *cmdTAC* under its native promoter. However, the ectopic production of Gp23 alone leads to protein aggregation and cellular toxicity as Gp23 requires an additional T4-encoded co-chaperonin, called Gp31, to fold properly^30,31^ (Fig. S3C). We therefore tested whether co-expressing *gp23* and *gp31* was sufficient to activate CmdT. Indeed, co-producing both Gp23 and Gp31 strongly inhibited growth in *cmdTAC*-containing cells, but not in an empty vector control (Fig. 3G). Importantly, Gp31 alone does not result in CmdT-dependent growth inhibition. Collectively, these results strongly indicated that Gp23 activates the CmdTAC system.

### CmdT is an mRNA ADP-ribosyltransferase activated in response to phage infection

How does CmdT lead to an abortive infection after its activation by Gp23 following phage infection? As noted earlier, sequence homology and structural predictions indicated that the CmdT toxin is likely an ADP-ribosyltransferase. To test for ADP-ribosylation, we first harvested protein from cells producing CmdTA for 60 minutes and blotted with an ADP-ribose specific antibody (Fig. 4A). The same bands appeared for cells producing CmdTA or harboring an empty vector indicating that CmdT was not modifying any cellular proteins. We also immunoblotted proteins extracted from cells harboring CmdTAC or an empty vector, each infected with T4 for 32 minutes (T4 replicates in ∼40 minutes in *E. coli* in these growth conditions). Again, the banding pattern was nearly identical in each case indicating that CmdT was not modifying proteins during phage infection (Fig. 4A). The bands seen following phage infection likely reflect the activity of known T4 ADP-ribosyltransferases such as Alt and ModB^32^.

**Figure 4.**
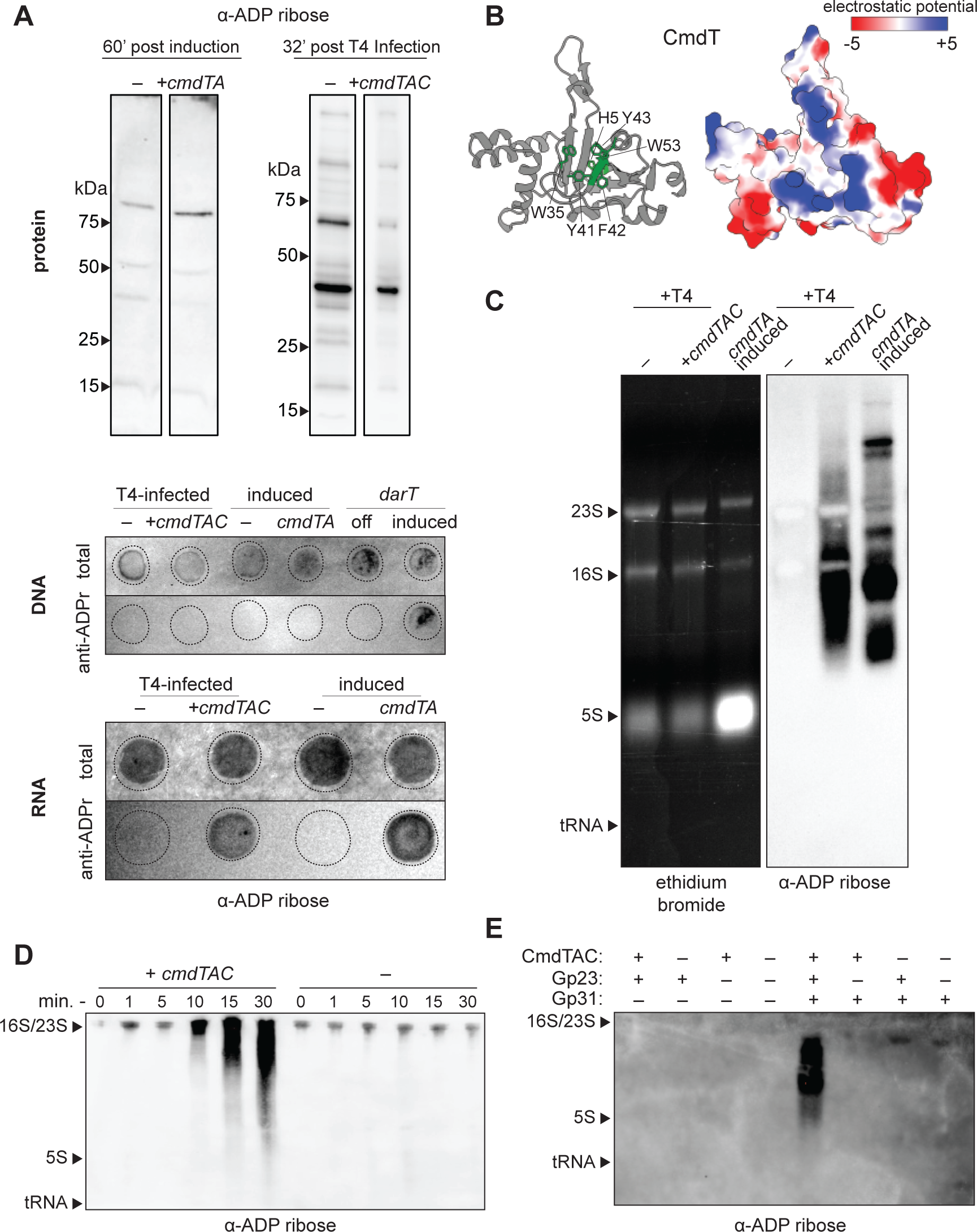
CmdT is an ADP-ribosyltransferase that specifically targets mRNA. (A) An anti-ADP-ribose antibody was used to probe protein gels (top) or DNA and RNA dot blots (bottom), as indicated. (Top, left) Samples taken from cells expressing *cmdTA* or harboring an empty vector, 60 minutes post-induction. (Top, right) Samples taken from cells expressing *cmdTAC* or harboring an empty vector and infected with T4 phage for 32 minutes. Proteins were separated by SDS-PAGE, while DNA and RNA samples were spotted onto nylon membranes. Total = methylene blue stained membranes. (B) AlphaFold2-predicted structure of CmdT (left) with conserved aromatic residues and putative catalytic residues (see Fig. 1G) shown in green, with the corresponding electrostatic surface representation (right). (C) RNA samples from strains and conditions indicated were resolved on agarose gels and then stained with ethidium bromide (EtBr) to visualize total RNA or blotted with an anti-ADP ribose antibody to visualize ADP-ribosylated RNAs. (D) Immuno-northern blotting of RNA samples taken from cells harboring *cmdTAC* or an empty vector at the times indicated post-infection with T4 phage. RNA was resolved on polyacrylamide gels. (E) Same as panel (D) but for cells harboring *cmdTAC*, or an empty vector, and producing the combinations of Gp23 and Gp31 indicated. RNA was resolved on polyacrylamide gels.

An electrostatic model of the predicted CmdT structure has a large positive cleft surrounding the region containing the putative catalytic and conserved aromatic residues (Fig. 4B). Given this positive patch, we hypothesized that CmdT might be binding to nucleic acids. To test for DNA modifications, we isolated DNA from cells harboring *cmdTAC* or an empty vector and infected with T4, as above, and performed a dot blot with an anti-ADP ribose antibody. However, there was no signal in either case indicating that CmdT does not modify DNA (Fig. 4A). Similarly, induction of *cmdTA* showed no signal, whereas inducing the expression of the DNA-targeting ADP-ribosyltransferase, DarT, did^10^. We then repeated the same procedure instead using total RNA. In this case, we observed strong signal on the anti-ADP ribose blot for cells harboring *cmdTAC* and infected with T4 or for cells expressing *cmdTA*, (Fig. 4A), but not in the empty vector controls, suggesting that CmdT was ADP-ribosylating RNA.

To better determine which RNA species were being modified, we extracted RNA from T4-infected cells harboring *cmdTAC* or an empty vector, or uninfected cells expressing *cmdTA* and ran them on an agarose gel for better resolution of higher molecular weight RNAs. We then transferred to nylon and probed with the anti-ADP ribose antibody (Fig. 4C). ADP-ribose signal was only seen for the *cmdTAC* samples. Strikingly, there was no signal corresponding to rRNA or tRNA, indicating that CmdT is likely an mRNA-specific ADP-ribosyltransferase. To the best of our knowledge, such an activity has not been reported before in bacteria or eukaryotes.

We then used ADP-ribosylation of mRNA to assess when, during T4 infection, CmdT is activated. Probing total RNA harvested at multiple timepoints over the course of T4 infection, we observed ADP-ribosylation of RNA within 10 minutes post-infection (Fig. 4D and Fig. S4A). We also tested whether inducing Gp23 and Gp31 in cells harboring CmdTAC, which leads to growth inhibition, led to ADP-ribosylation of RNA. We tested the expression of all combinations of *cmdTAC*, *gp23*, and *gp31*, and found that only when all three were expressed could we detect RNA modification (Fig. 4E and Fig. S4B). Collectively, our results indicate that CmdT is an RNA-targeting ADP-ribosyltransferase activated by the presence of T4 Gp23 and Gp31.

### CmdT blocks translation and the expression of late genes to prevent T4 replication

To understand how mRNA ADP-ribosylation inhibits phage development, we measured translation by growing T4-infected cells producing CmdTAC or CmdT*AC in the presence of radiolabeled cysteine and methionine. By 10 minutes post-infection we observed a complete block in translation in cells with CmdTAC but not CmdT*AC (Fig. 5A). The rapid block in translation was consistent with our observation of ADP-ribosylated mRNA on immuno-Northern blots 10 minutes post-infection (Fig. 4D).

**Figure 5.**
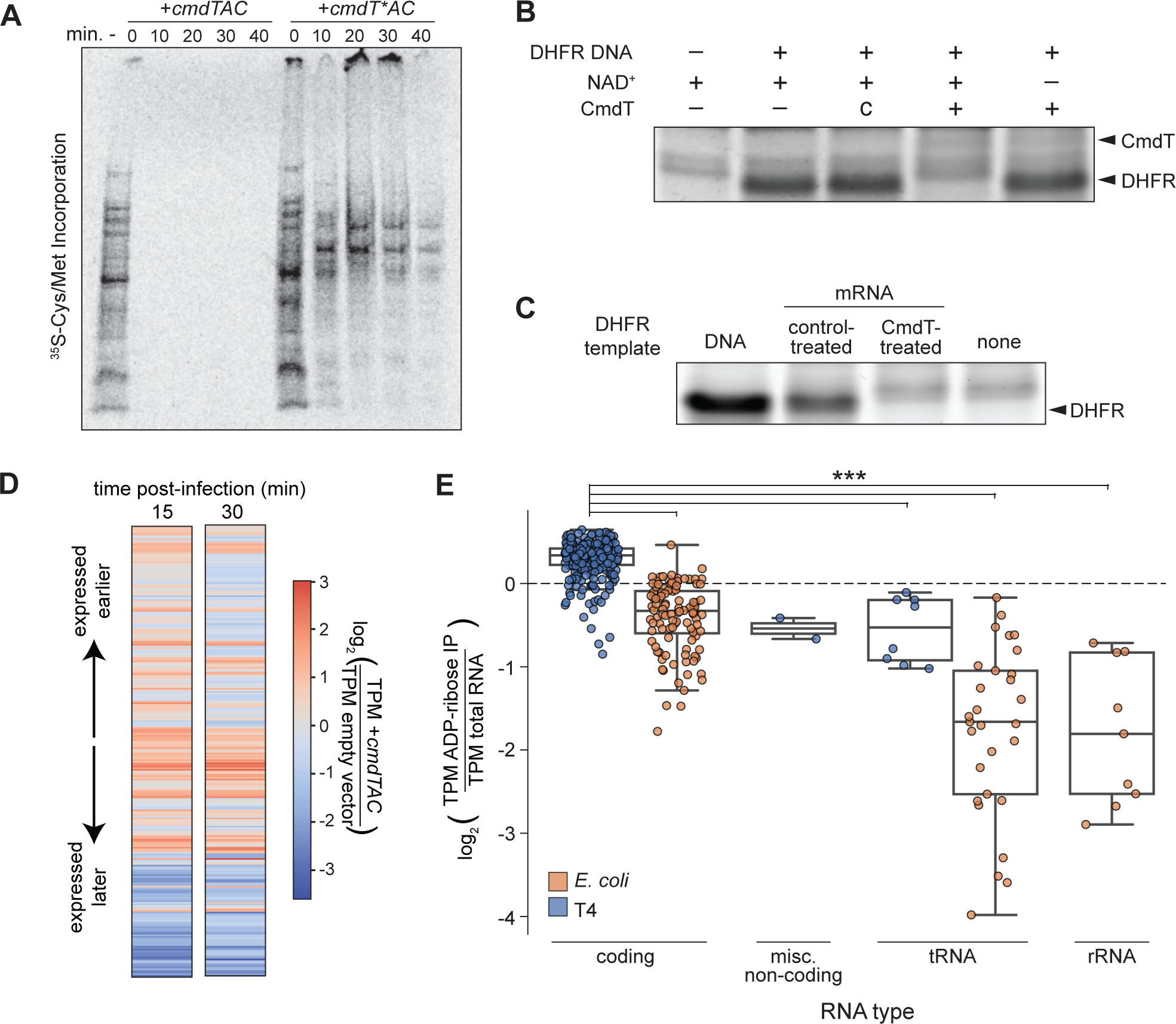
CmdT blocks translation and the transition to late-stage T4 infection. (A) Protein synthesis, as measured by ^35^S-labeled cysteine and methionine incorporation at the time points indicated during T4 infection of cells harboring CmdTAC or a variant, CmdT*AC, with a catalytically inactive CmdT. Proteins were resolved by SDS-PAGE before phosphorimaging. Data shown are a representative image from two independent biological replicates. (B) *In vitro* transcription-translation reactions with DNA encoding dihydrofolate reductase (DHFR) used as template. Purified CmdT and NAD^+^ were added, where indicated, with DHFR production visualized by Coomassie staining, c = control eluant. (C) *In vitro* transcription-translation reactions using a DNA template encoding DHFR or mRNA templates produced by T7 RNA polymerase and treated with CmdT (or eluent from a mock purification of untagged CmdT) prior to being added. (D) Heat maps summarizing, for each T4 transcript, the ratio of TPM values in cells harboring *cmdTAC* or an empty vector. Transcripts are ordered from top to bottom based on peak time of expression, as determined previously^15^. Data are the average of two biological replicates. (E) Ratio of ADP-ribose IP enriched TPM to baseline RNA TPM values for each transcript. Transcripts are separated by RNA type and T4 vs MG1655 origin. Note that the T4 genome does not contain rRNA. Asterisks indicate p-value < 0.01 for a Welch’s t-test. Data are the average of two biological replicates.

To further test if CmdT activity blocks translation, we purified CmdT-His_6_, and, as a control, we concentrated protein eluted from a mock purification of CmdT lacking an affinity tag. Purified CmdT, but not the control eluant, blocked translation of a model protein, DHFR, in a cell-free *in vitro* transcription/translation reaction, and inhibition was dependent on NAD^+^ (Fig. 5B). To ensure that CmdT had blocked DHFR translation, not transcription, we first *in vitro* transcribed DHFR and then treated the mRNA with purified CmdT and NAD^+^. Providing this template to the *in vitro* translation reaction again resulted in no DHFR protein. In contrast, purified DHFR mRNA treated with the control eluant was robustly translated (Fig. 5C). Together, these data suggest that CmdT modifies mRNA to block translation.

To understand the global effect of CmdT activation on the T4 lifecycle *in vivo,* we performed RNA-seq during phage infection. We harvested RNA at 15- and 30-minutes post-infection of both CmdTAC-containing and control cells, generated RNA-seq libraries, and then mapped the reads back to the T4 genome. In comparison to the empty vector control, T4 infected *cmdTAC+* cells showed substantially decreased expression of late genes and, consequently, increased read counts from early and middle genes; the loss of late transcripts increases the relative percentage of middle and early transcripts (Fig. 5D). The various stages of T4 gene expression depend on the successful translation of gene products in earlier stages of the lifecycle^33^ suggesting that T4 replicating in *cmdTAC+* cells are unable to progress to the late stages. This conclusion is consistent with our finding that CmdT-dependent ADP-ribosylation of mRNA can be detected within 10 minutes post-infection and that CmdT almost completely blocks translation.

We also performed an RNA-immunoprecipitation with sequencing (RIP-seq) experiment by enriching for ADP-ribosylated RNAs with an anti-ADP-ribose specific antibody, followed by deep sequencing. We harvested RNA from *cmdTAC+* cells at 30 minutes post-infection, and then subjected this RNA to anti-ADP-ribose immunoprecipitation before generating an RNA-seq library. In addition to the immunoprecipitated RNA, we also generated traditional RNA-seq libraries from the same harvested RNA to assess transcript coverage. Transcripts per million reads (TPM) values for T4 mRNAs were significantly higher in the IP samples over the control RNA-seq (p < 0.05; Welch’s t-test). We then calculated a ratio of TPM values in the IP and control samples. These ratios were significantly higher for T4 mRNAs than for tRNAs and rRNAs (Fig. 5E). These findings reinforce our conclusion that CmdT is mRNA specific. Based on our findings, we conclude that the ADP-ribosylation of mRNA by CmdT leads to a halt in translation, preventing the progression of T4 into the late stages of its life cycle and a failure to produce mature progeny.

### CmdT specifically modifies GA dinucleotides in single-stranded RNA

To determine the specificity of nucleic acid modification by CmdT we first performed *in vitro* ADP-ribosylation reactions by mixing purified CmdT, NAD^+^, and a 24 nt model RNA oligo which contained a Shine-Dalgarno sequence and an AUG start codon. We found that CmdT treatment resulted in a shift in the migration of the ssRNA substrate on denaturing gels, indicative of covalent RNA modification (Fig. 6A). This shift was dependent on the inclusion of NAD^+^ in the reaction (Fig. 6A). When a ssDNA oligo with the same sequence was supplied as a substrate, a product was formed, but with substantially reduced kinetics (Fig. 6B). When we supplied the equivalent double stranded RNA oligo as a substrate, we did not observe a product, even at high concentrations of CmdT (Fig. 6C).

**Figure 6.**
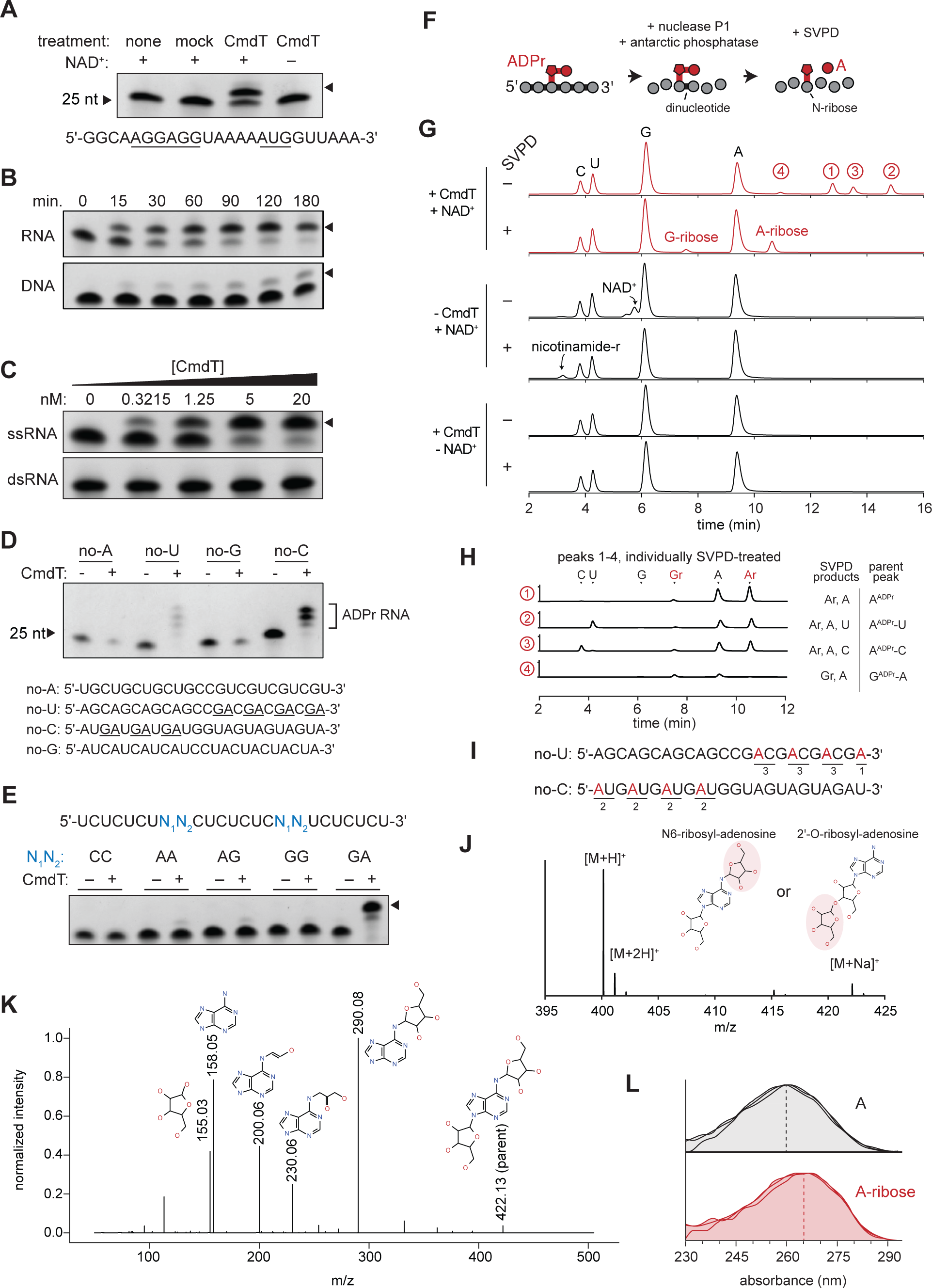
CmdT modifies the N6 methyl group of adenine in GA dinucleotides of single-stranded RNA. (A) The RNA oligo indicated was incubated with ∼ 4 nM CmdT-His_6_, protein from a mock purification of untagged CmdT, or no protein, along with NAD^+^ where indicated. Reaction products were resolved on a polyacrylamide gel and imaged by methylene blue staining. (B) ADP-ribosylation by ∼ 4 nM CmdT-His_6_ of the RNA (top) or equivalent DNA oligo (bottom) at the time points indicated, visualized as in (A). (C) ADP-ribosylation by CmdT-His_6_ of the ssRNA oligo from (B) or the corresponding dsRNA at increasing concentrations of CmdT-His_6_ for 60 min. (D) ADP-ribosylation by CmdT-His_6_ of the oligos listed at the bottom containing no A, U, G, or C nucleotides. (E) ADP-ribosylation by CmdT-His_6_ of the oligo indicated (top) with the identity of the variable dinucleotides indicated above each reaction. (F) Schematic showing enzymatic digestion of ADP-ribosylated oligos with nuclease PI and antarctic phosphatase, with or without snake venom phosphodiesterase (SVPD). (G) HPLC analysis of nucleosides isolated following incubation of the no-U and no-C RNA oligos in panel (D) with CmdT and NAD^+^, as indicated, and then treated with nuclease P1 and antarctic phosphatase, and snake venom phosphodiesterase (SVPD), where indicated. Peaks corresponding to A, C, G, and U nucleosides, NAD^+^, G-ribose, and A-ribose, and nicotinamide-riboside (labeled nicotinamide-r; produced from SVPD cleavage of unused NAD^+^) are marked. The y-axis represents milli-absorbance units (mAu) at 254 nm. ADPr = ADP-ribose. (H-I) HPLC analysis of nucleosides produced after treating the numbered peaks from panel (F) with SVPD. The products are labeled and the inferred parent molecule indicated on the far right with the origin of the peaks on the oligo substrates shown in (H). Gr = G-ribose; Ar = A-ribose; ADPr = ADP-ribose. (J) ESI MS analysis of the A-ribose peak from panel (F), structures shown with implicit hydrogens. (K) MS/MS fragmentation of the A-ribose sodium adduct. Collision energy, 45 eV. Predicted fragments are annotated and m/z is indicated. (L) UV-Vis spectroscopy of adenosine and CmdT-produced adenosine-ribose from (F).

We also incubated purified CmdT with total RNA extracted from cells infected with T4, supplying 6-biotin-17-NAD^+^ as a substrate for ADP-ribosylation. We then enriched for ADP-ribosylated RNA by precipitation with streptavidin beads and sequenced RNA samples taken pre- and post-streptavidin pulldown, without rRNA depletion (Fig. S5). The results of this experiment paralleled those of the *in vivo* RIP-seq, with enrichment of mRNAs compared to tRNAs and rRNAs. Together, these data suggested that unstructured, ssRNA is the preferred substrate of CmdT ADP-ribosylation.

Next, we attempted to determine which nucleotide base is modified, and whether CmdT targets specific sequence motifs. We repeated the ADP ribosylation reactions with CmdT and NAD^+^ incubated with one of four ssRNA oligo substrates, each missing one base. Only the no-U and no-C oligos produced slower migrating products, suggesting that neither C nor U is required for modification (Fig. 6D). Notably, the no-U and no-C substrates produced four and three products, respectively, each of decreasing mobility in the gel (Fig. 6D, Fig. S6A). This observation suggested that CmdT was adding multiple modifications to individual oligos. The formation of the slower migrating bands was also dependent on reaction time (Fig. S6A). We noticed that the number of bands formed equated with the number of 5’-AG-3’ or 5’-GA-3’ dinucleotides in the no-U and no-C oligos (Fig. 6D). Similarly, the model RNA used in Fig. 6A-6C contains one GA site, and produced one reaction product. These observations indicated that CmdT requires GA dinucleotides to catalyze ADP ribosylation.

To further test the sequence specificity of CmdT, we designed five ssRNA oligos with invariant C- and U-containing scaffolds. At two separate positions within the oligos, we introduced various dinucleotides: CC (control), AA, AG, GG, or GA. When we supplied these oligos as substrates in ADP ribosylation reactions, substantial product was only produced with the substrate containing GA at the variable positions (Fig. 6E). Very faint bands appeared when the variable dinucleotide contained two purines other than GA, but none when the oligo contained two pyrimidines (CC). In addition, two products were formed from the GA oligo substrate, suggesting that the presence of two GA sites results in the formation of two modified ssRNA products. Taken together, our results indicate that CmdT preferentially ADP-ribosylates ssRNA and recognizes GA dinucleotides.

To determine which nucleotides are covalently modified with ADP-ribose, we performed a CmdT ADP-ribosylation reaction on an equimolar mixture of the no-U and no-C RNA oligos. We then digested the ADP-ribosylated RNA using nuclease P1, which hydrolyzes 3’-5’ phosphodiester bonds, and antarctic phosphatase, which removes 5’ and 3’ phosphates, and resolved the products with HPLC (Fig. 6F, also see Methods and Fig. S6C). In addition to peaks corresponding to C, U, A, and G nucleosides, we observed 4 other peaks (Fig. 6G, top panel). These additional peaks were not seen in control reactions lacking NAD^+^ or lacking CmdT. Notably, the absorbance of the adenosine peak dropped substantially in CmdT-treated samples compared to the control reactions, suggesting that the four new peaks were likely derived mainly from modified adenosine (Fig. S6B).

In parallel, we incubated the same reactions with snake venom phosphodiesterase I (SVPD), in addition to the nuclease and phosphatase. SVPD cleaves the phosphodiester bond in ADP-ribose, in addition to some phosphodiester bonds that may be inaccessible to nuclease P1 (Fig. 6F). For example, SVPD treatment of ADP-ribosylated adenosine would produce ribosylated adenosine and adenosine, with the ribose and adenosine originating from ADP-ribose (Fig. S6C). In this way, CmdT-targeted nucleosides can be detected as ribose-modified derivatives in HPLC. For the samples treated with CmdT and NAD^+^, SVPD treatment produced two peaks with relative retention times corresponding to adenosine-ribose and guanosine-ribose^11^, with the former showing stronger signal (Fig. 6G). Because SVPD treatment resolved four CmdT-dependent species to two, adenosine-ribose and guanosine-ribose, it is likely that at least some of the original four peaks are dinucleotides in which the phosphodiester bond between nucleotides is inaccessible to nuclease P1 but hydrolyzed by SVPD^11^.

To identify the four CmdT-dependent products observed before SVPD treatment (Fig. 6G, top panel), we isolated fractions corresponding to each peak, treated each with SVPD, and then re-ran HPLC. For fraction 1, this procedure produced two major peaks corresponding to adenosine-ribose and adenosine indicating that the parent peak was ADP-ribosylated adenosine (Fig. 6H). Treatment of fractions 2 and 3 produced the same two peaks, along with peaks of uridine and cytosine, respectively, indicating that the parent molecules were the dinucleotides AU and AC, with ADP-ribose on adenosine in each case (Fig. 6H-I). Finally, treatment of fraction 4 produced guanosine-ribose and adenosine peaks indicating the parent molecule was the dinucleotide GA with the guanosine ADP-ribosylated. Importantly, because the majority of each sample resolved to adenosine-ribose, we conclude that adenosine is the primary target of ADP-ribosylation, with occasional modification of guanosines, at GA, AG, or GG sites. Taken together, our results suggest that CmdT primarily ADP-ribosylates adenosine nucleotides at 5’-GA-3’ motifs in single-stranded mRNA.

To confirm the identity of the peak we inferred to be adenosine-ribose, we performed electrospray ionization mass spectrometry (ESI-MS), which revealed peaks at 400 and 422 m/z, equivalent to the molecular weight of adenosine-ribose [M+H]^+^ and [M+Na]^+^ (Fig. 6J). There are two positions on adenosine most likely to react with NAD^+^ and become ADP-ribosylated, the 2’ hydroxyl of the ribose, or N6 on the adenine base (Fig. 6J). To distinguish between these, we subjected the adenosine-ribose sodium adduct to ESI-MS/MS fragmentation analysis (Fig. 6K and Fig. S6D). We did not detect a fragment ion for ribose-2’-O-ribose (m/z 287) which should be produced if the 2’-OH is ADP-ribosylated^11^. Instead, we observed peaks at 200 and 230 m/z, which likely correspond to fragments in which the adenine base contains a portion of the ribose derived from ADP-ribosylation of N6 (Fig. S7).

To further test that the adenine base is modified, we compared the UV absorbance (λ_max_) of adenosine and CmdT-generated adenosine-ribose. ADP-ribosylation at the 2’ hydroxyl of ribose does not shift the UV absorbance spectrum of adenosine^11^, while modifications of the N6 position, such as N6-methylation or N6-isopentenylation, redshift the UV absorbance of adenosine by 6 or 8 nm, respectively^34^. The adenosine-ribose produced by CmdT shifted the λ_max_ from 257 nm to ∼265 nm (Fig. 6L). Given our mass spectrometry and UV spectroscopy data, along with the finding that CmdT can weakly modify ssDNA (which lacks a 2’-OH), we conclude that the site of ADP-ribosylation by CmdT is the N6 of adenine in GA dinucleotides within single-stranded RNA.

## Discussion

### ADP-ribosylation of mRNA

Our work reveals the first case, to our knowledge, of an ADP-ribosyltransferase that specifically targets mRNA. Prior studies have identified numerous ADP-ribosyltransferases in phages, bacteria, and eukaryotes that use the highly abundant and reactive molecule NAD^+^ to covalently modify a wide range of proteins, often to reversibly regulate their activities^1–3^. Protein ADP-ribosyltransferases are also featured in many biological conflicts with secreted toxins such as pertussis and cholix toxins capable of shutting down key cellular processes by modifying specific target proteins^5,35^. More recently, ADP-ribosyltransferases that target DNA and structured RNAs have been found^8,11,12^. Our findings extend the range of targets to include mRNA, with CmdT-catalyzed ADP-ribosylation leading to a potent block in protein translation. Notably, *in vitro* studies have shown that human PARP10, PARP11, PARP14, and PARP15 can each modify single-stranded RNA substrates^7,36^, though evidence for RNA modification *in vivo* is so far lacking. Nevertheless, these prior results suggest that ADP-ribosylation of RNA may be a widespread, but previously unappreciated modification occurring throughout biology. PARP10, PARP11, and PARP14 are also known interferon-stimulated genes^37–39^ and several mammalian viruses, including SARS-Cov2, encode ADP-ribosylhydrolases that offset PARP activity to promote viral replication^40,41^. Thus, the ADP-ribosylation of viral RNAs may be a broadly conserved and critical facet of anti-viral immunity.

Our results indicate that CmdT primarily modifies the N6 position of adenines within GA dinucleotides. Relying on such a short motif for modification likely provides CmdT with two key advantages. First, GA nucleotides are almost always found with the Shine-Dalgarno (SD) sequences of bacterial and phage transcripts. *E. coli* transcripts typically have a SD sequence matching or similar to the consensus AGGAGG^42^, whereas T4 early genes rely on the slightly shorter GAGG^33^. ADP-ribosylation of SD sequences may prevent modified transcripts from engaging ribosomes and initiating translation. Alternatively, or in addition, the modification of GA dinucleotides within an mRNA may disrupt base-pairing with tRNAs or may stall ribosomal translocation. Further investigation is required to understand exactly how the block to translation occurs. Second, unlike restriction-modification systems or CRISPR-Cas, which target 4-32 bp motifs, it would be implausible for a phage to escape CmdTAC by acquiring mutations to its recognition sites in a phage genome^43^.

The structural basis of CmdT’s ability to ADP-ribosylate GA dinucleotides will also require future work. However, we noted that the AlphaFold2-predicted structure of CmdT has an extensive electropositive region on its surface that likely facilitates interaction with its negatively charged RNA substrates. We also noted that CmdT homologs contain four highly conserved aromatic residues that are not part of the catalytic triad (Fig. 1G). These residues may facilitate τ-stacking with the purine rings of a GA dinucleotide targeted by CmdT^44^.

### Direct triggers of anti-phage defense

Dozens of new anti-phage defense systems have been discovered recently^21,45,46^, most of which function through an Abi mechanism in which infected cells kill themselves or stop growing to prevent the production of new phages. Abi-based systems must remain off in the absence of infection and then be rapidly triggered upon phage infection. But how Abi systems sense phage infection remains poorly understood. Our work reveals the T4 major capsid protein as the likely direct activator of the CmdTAC system (Fig. 7). Structural proteins appear to be common triggers of anti-phage defense systems, with prior work revealing the major capsid protein of SECβ27 as the trigger of CapRel^SJ46^ (ref^16^) and terminase and portal proteins as the direct activators of Avs1-3 and Avs4, respectively^47^. Based on IP-MS, we found that the T4 major capsid protein Gp23 physically associated with the CmdC chaperone, and was sufficient, when co-produced with the phage-encoded chaperonin Gp31, to liberate CmdT from CmdAC in the absence of phage infection. *Gp23* is classified as a late gene based on its time of maximal expression, but can be detected earlier in RNA-seq and proteomic studies^15,48^. We detected the CmdC-Gp23 interaction by IP-MS at 15 minutes post-infection, and our immunoblotting studies indicated that ADP-ribosylated RNA could be detected as early as 10 minutes post-infection, supporting the notion that enough Gp23 is produced at that stage to trigger CmdT activity.

**Figure 7.**
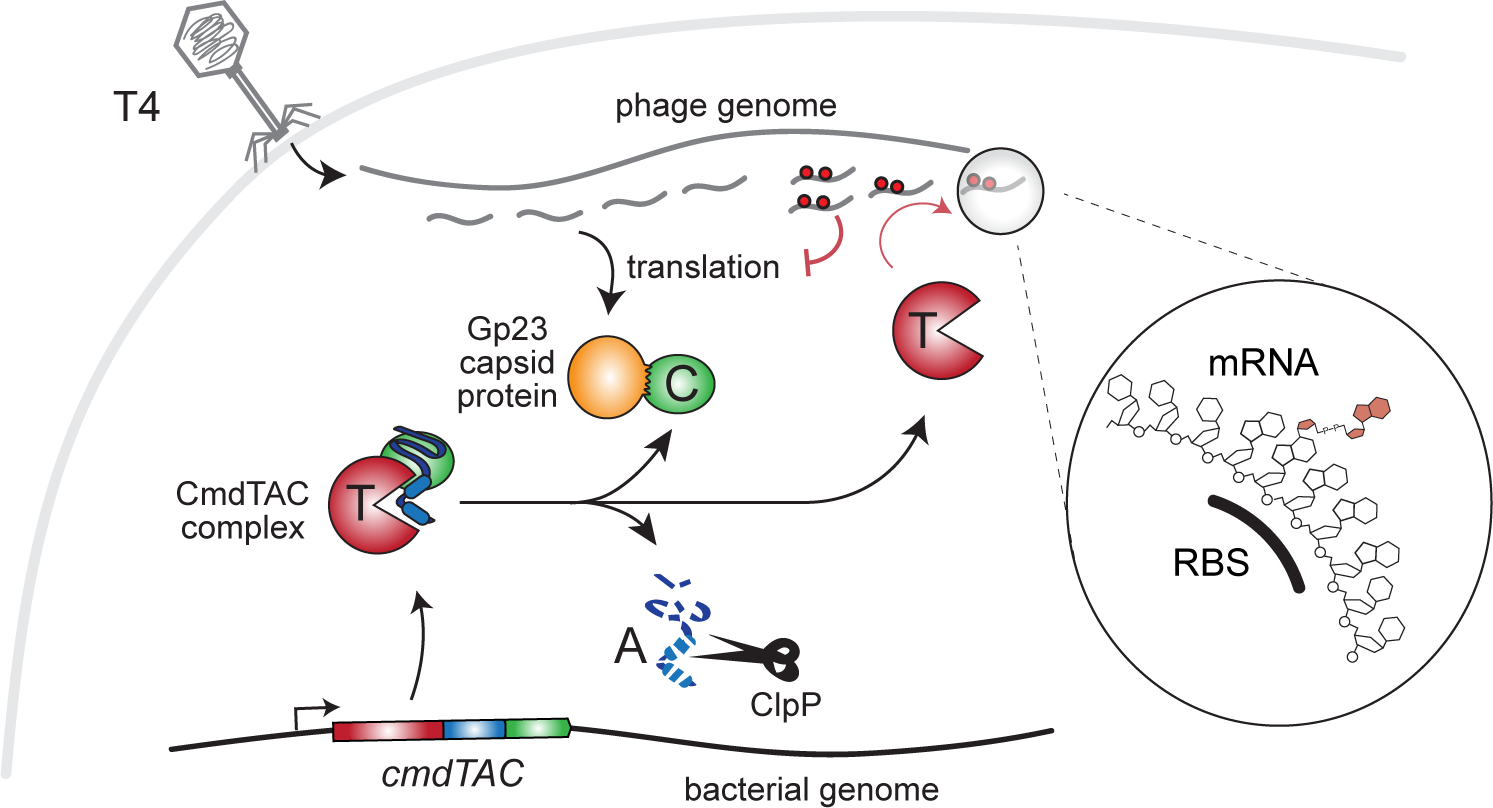
Model for anti-phage defense by the CmdTAC system. CmdTAC produced by *E. coli* forms a complex prior to infection. Following T4 infection, newly synthesized capsid protein (Gp23) binds CmdC chaperone, leading to degradation of CmdA by ClpP and subsequent release of active CmdT toxin. CmdT ADP-ribosylates GA dinucleotides in single-stranded RNAs, including within the Shine-Dalgarno sequence of most T4 transcripts, thereby preventing their translation and the production of mature T4 virions.

Precisely how the capsid protein interacts with CmdC and the specific regions involved will require further biochemical studies. Notably, despite extensive directed evolution we did not isolate any escape mutants of *gp23*, an essential gene. This could indicate that mutations to *gp23* that escape binding to CmdC are lethal by preventing the production of stable, mature virions. In addition, CmdC may engage an extended region of the capsid protein such that multiple substitutions and complex mutations would be required to abrogate binding.

Phage structural proteins may be common triggers for anti-phage defense systems as they typically adopt folds that differ substantially from endogenous, host proteins, thereby providing requisite specificity. Additionally, structural proteins must accumulate to high levels, helping to ensure defense system activation. One potential downside to relying on structural proteins as triggers is that most are encoded by late genes, raising the possibility that activation could fail to happen before new virions are produced. However, the ‘late’ classification is based on the peak time of expression and both RNA-seq and mass spectrometry studies indicate that late genes are expressed at earlier time points before peaking later in the phage’s life cycle^15,48^. Consistent with this idea, we detected CmdT activation and the ADP-ribosylation of mRNAs as early as 10 minutes post-infection.

### TAC systems in anti-phage defense

CmdTAC adds to the growing list of TA systems shown to function in anti-phage defense^13^ and represents the first such TAC system. TAC systems have been relatively well-characterized outside the context of phages^17–20^, revealing common principles such as the reliance of their antitoxins on C-terminal chaperone-addiction domains for folding and stability, and the use of SecB-like chaperones. The toxins of TAC systems show more variability, including several different families of endoribonucleases, which could, in principle, function like CmdT to inhibit translation and prevent phage replication. More broadly, our discovery of ADP-ribosyltransferase activity in CmdT underscores the tremendous diversity in the mechanisms of action of toxins associated with TA and TAC systems. Many of these may act similarly to CmdTAC and other recently characterized anti-phage defense systems, inhibiting a range of key cellular processes to either kill or block the growth of phage-infected cells to prevent the production of mature virions.

As with virtually all anti-phage defense systems discovered to date, phages have likely evolved mechanisms to counteract the CmdTAC system. We isolated mutations in T4 Alt.-3 – a protein of unknown function – that abolished protection by CmdTAC by blocking degradation of CmdA, preventing CmdT liberation. In principle, phages could have native proteins that inhibit the ClpP protease to stabilize CmdA and thereby indirectly inhibit CmdT activity. Indeed, T4 has been known for decades to produce the Lon protease inhibitor PinA^49^, so it’s conceivable that ClpP inhibitors are produced by other phages as a means of inhibiting TA-or TAC-based defense systems. Phages could also produce SecB-like chaperones that stabilize the CmdTA complex to prevent CmdT liberation. Counter-defense to CmdT may also involve direct inhibition, as seen with other defensive TA systems like Dmd, which inhibits the RnlAB system^50^, or indirect mechanisms such as an ADP-ribosylhydrolase that can ‘undo’ the modifications made by CmdT.

### Concluding Remarks

The intense conflict between phages and the bacteria they prey upon has led to the evolution of sophisticated anti-phage defense systems with a wide range of molecular innovations. Many of these systems have been identified in recent years, but the molecular mechanisms by which they inhibit phages remain incompletely defined. Our work now reveals mRNA ADP-ribosylation as a potent defense mechanism used by bacteria to thwart phages. As several interferon-inducible mammalian PARPs – whose exact functions remain unclear – can also modify RNA, this mechanism may be broadly used in immunity throughout biology.

## Acknowledgments

We thank D. Saxton, T. Zhang, S. Srikant, and C. Beck for comments on the manuscript, B. Imperiali for help interpreting the mass spectrometry data, C. Eickmann for help with AlphaFold2 predictions, the MIT BioMicro center and its staff for their support with sequencing, and the MIT Biopolymers and Proteomics Core and its staff for assisting in HPLC and MS experiments. We also thank S. Srikant for help with T4 reverse genetics. M.T.L. is an Investigator of the Howard Hughes Medical Institute. This work was also supported by a Howard Hughes Medical Institute Gilliam Fellowship awarded to C.D and a National Institutes of Health NIGMS grant 5F32 GM139231-02 to C.N.V.

## Author Contributions

C.R.D. and C.N.V. performed all experiments. C.R.D., C.N.V., and M.T.L. designed experiments, analyzed data, prepared figures, and wrote the manuscript.

## Declaration of Interests

The authors declare no competing interests.

## Methods

### Data and Code Availability

Raw data for IP-MS/MS of CmdC pulldown can be accessed at doi:10.25345/C52J68G0H. Raw data for nucleotide MS and ESI-MS/MS can be accessed at doi:10.25345/C51N7XZ2Q. RNA-seq and RIP-seq data are available at GEO under accession number GSE253514. Code used for RNA-seq and *in* vivo RIP-seq analysis is available at doi:10.5281/zenodo.10522487.

### Bacterial and phage growth and culture conditions

1. *E. coli* was routinely grown in LB medium at 37 °C unless otherwise stated. Phages were propagated and handled as described previously^21^.

### Strain construction

All primers and gBlocks used in plasmid construction can be found in Table S1. All plasmids and strains are listed in Tables S2-S3. gBlocks were synthesized by IDT. Strains were typically constructed by PCR amplifying vector and insert fragments with compatible 25-30 bp overhangs and ligating them by Gibson assembly (NEB). For the *alt.-3*-targeting pCas9 construct, first, the spacer oligonucleotides were ligated in 50 mM NaCl by heating to 98 °C and slowly cooling to 25 °C over 30 minutes. Then Golden Gate Assembly was conducted with PCR amplified vector, annealed oligos, T4 DNA ligase (NEB) and BsaI-v2 (NEB). Site-directed mutagenesis (Y41A) and insertion of epitope tags was accomplished by PCR amplifying vector template for 15 cycles with two, complementary ∼60 bp oligonucleotides containing the target mutation or insertion.

### Plaque assays

Overnight cultures of indicated strains were mixed 1:80 with melted 0.5% LB agar and then overlaid on 1.2% LB agar plates. For plaque assays done with induction of an arabinose-inducible promoter, base layer plates contained 0.2% w/w arabinose. A 10-fold dilution series of the indicated phage was spotted onto plates and the plates grown at 30 °C overnight and plaque-forming units (PFU) were enumerated. Log_10_ protection (Fig. 1C) was measured as −log_10_ EOP where EOP is the ratio of PFU_experimental_/PFU_control_.

### Growth curves

For measuring growth during T4 infection, overnight cultures of *+cmdTAC* and *-cmdTAC* cells were back-diluted 1:200 in 96-well plates and infected with T4 at the indicated MOIs. Cultures were grown at 37 °C with orbital shaking on a plate reader (Biotek) for 6 hours. For ectopic expression of Gp23 and Gp31 with CmdTAC, overnight cultures were back-diluted to OD_600_ of 0.05 in M9L + 0.2% w/w glucose + 100 ng/mL aTc and grown for 3 hours at 37 °C to pre-induce Gp31. Cultures were then pelleted and resuspended at an OD_600_ of 0.05 in fresh M9L + 0.2% w/w glucose + 100 ng/mL aTc or M9L + 0.2% w/w glucose + 100 ng/mL aTc. Cultures were grown at 37 °C with orbital shaking on a plate reader for 12 hours.

### RNA extraction following phage infection

Overnight cultures of *+cmdTAC* and *-cmdTAC* cells were back-diluted and grown at 37 °C to OD_600_ between 0.2 and 0.3 before being infected with T4 at a MOI of 10. RNA was extracted from cells at multiple timepoints post-infection as previously described^15^. Briefly, 1 mL of cells was mixed with 1 mL of boiling lysis buffer (SDS 2%, 4 mM EDTA pH = 8) and incubated at 100 °C for 5 min before flash freezing in liquid nitrogen. 2 mL of acid-buffered phenol solution (pH 6, Sigma) heated to 67 °C was added to thawed samples, vortexed, and then incubated at 67 °C for 2 minutes. Samples were spun down at 20,000 *g* for 10 minutes and hot phenol extraction repeated on the collected aqueous layer. A third extraction was then done using 2 mL of acid-buffered phenol-chloroform solution (Ambion). RNA from the final extraction was then precipitated at −20 °C for at least 1 hour or at −80 °C overnight with 1x volume isopropanol, 1/10x volume 3M NaOAc (pH 5.5, Thermo Fisher), and 1/100x volume GlycoBlue. RNA was pelleted via centrifugation at 4 °C and 20,000 *g* for 30 min. Pellets were washed twice with 800 mL of ice-cold 70% ethanol, air-dried, and resuspended in 90 μL RNAse-free H_2_O (Thermo Fisher).

To remove DNA, 10 μL of 10X Turbo DNAse buffer (Ambion) and 2 μL of Turbo DNAse I (Ambion) was added to each sample and incubated at 37 °C for 20 minutes. An additional 2 μL of Turbo DNAse I was then added, and samples again incubated at 37 °C for 20 minutes. RNA was extracted from this digest by precipitation with 3x volume ethanol, 1/10x volume 3M NaOAc (pH 5.5), and 1/100x volume GlycoBlue. Pelleting and washing were performed the same as described above. RNA yield was verified using a NanoDrop spectrophotometer.

### RNA extraction from non-infected cells

Cells were grown until desired conditions and then 900 μL of culture was mixed with 100 μL of stop solution (5% acid phenol, 95% ethanol) and inverted to mix. Samples were then spun down at 13,000 *g* for 30 seconds, the supernatant removed, and pellets flash frozen in liquid nitrogen. To each pellet, 400 μL of TRIzol Reagent (Invitrogen) heated to 65 °C was added and mixed using a thermomixer for 10 minutes at 65 °C and 2000 rpm before freezing at −80 °C for at least 10 minutes. Samples were thawed and then centrifuged at 20,000 *g* for 5 minutes at 4 °C to pellet any debris and the TRIzol solution moved to a new tube. RNA was purified using the Direct-zol RNA Miniprep kit (Zymo Research) following manufacture’s protocol including optional on-column DNAse treatment. RNA yield was verified using a NanoDrop spectrophotometer.

### Immuno-northern blotting

Novex 6% TBE-urea gels in 1x TBE buffer (Invitrogen) were pre-run at 180 V for at least 50 minutes prior to sample loading. Each RNA sample was mixed with equal volume of Novex 2X TBE-urea sample buffer (Invitrogen), heated at 90 °C for 10 minutes, and then placed on ice for 2-3 minutes just before loading. Gels were run at 180 V for 30-50 minutes depending on expected product length. Gels were removed from casing and incubated in 40 mL 1x TBE with added 4 μL of SYBR Gold stain (Thermo Fischer) for 10 minutes. Gels were imaged on a ChemiDoc MP imaging system (Bio-Rad) set for SYBR Gold imaging. RNA was transferred from the gel to a Hybond-N^+^ nylon membrane (Cytiva) via semi-dry transfer at 0.38 A for 90 minutes. After transfer, RNA was bound to the membrane by exposure to 120,000 μJ of UV radiation in a Stratalinker UV Crosslinker. Membranes were then incubated with shaking in 0.2% iBlock (Invitrogen) in 1x PBST for 10 minutes at room temperature or overnight at 4 °C. Primary antibody treatment was done with Poly/Mono-ADP Ribose rabbit antibody (Cell Signaling Technologies) diluted 1:1000 in 0.2% iBlock + 1x PBST either for 2 hours at room temperature or overnight at 4 °C with shaking. Following primary antibody treatment, membranes were washed 3 times for 10 minutes each with 1x PBST. For secondary antibody treatment, membranes were incubated for 1 hour with shaking at room temperature with goat anti-rabbit IgG (H+L) secondary antibody, HRP (Invitrogen) diluted 1:1000 in 0.2% iBlock + 1x PBST. Membranes were then again washed 3 times for 10 minutes each in 1x PBST. Signal was developed using SuperSignal West Femto maximum sensitivity substrate (Thermo Fischer) and imaged on a ChemiDoc MP imaging system set for chemiluminescence detection. Dot blots were conducted identically except 250 ng DNA or 1 µg RNA were spotted on membranes.

For agarose immuno-northern blots, 0.8 g of agarose was melted in 66.7 mL of H_2_O and allowed to cool to 65 °C. 8 mL of 0.2 M (10x) 3-(N-morpholino)propanesulfonic acid (MOPS) buffer, 5.4 mL of formaldehyde and 5 µl of 10 mg/mL ethidium bromide (EtBr) were added to the agarose, and a 14−12 cm gel was cast and allowed to cool. 4 µg RNA were added to 17 µL sample buffer (2 µL 10X MOPS, 4 µL formaldehyde, 10 µL de-ionized formamide, and 1 µL EtBr) and samples were denatured at 80 °C for 10 minutes then cooled on ice for 5 minutes. Prior to sample loading, the empty gel was run at 115 V for 5 minutes. 2 µL of loading dye (50% glycerol, bromophenol blue, and xylene cyanol) were added to each RNA sample. Samples were then electrophoresed at 100V for 80 minutes in 1x MOPS buffer. The gel was visualized before soaking in H_2_O for 10 minutes followed by a 20-minute equilibration in transfer buffer (3 M NaCl, 0.01 N NaOH). RNA was transferred onto Hybond-N+ nylon membrane by upward capillary transfer at room temperature for 75 minutes in transfer buffer. Immunoblotting was performed as described above.

### RNA immunoprecipitation and sequencing

Cells were collected and RNA harvested as described above for infected cells. rRNA was removed using a previously described ribosomal RNA subtraction method^68^. rRNA-depleted RNA was then fragmented using sonication. For each sample to be sonicated, 4 μg of RNA was added to 100 μL 1x TE buffer (Sigma) in a 1.5 mL TPX microtube (Diagenode) and incubated on ice for 15 minutes. Tubes were then placed in a Bioruptor 300 sonicator water bath chilled to 4 °C for 60 cycles of 30 seconds on, 30 seconds off at high power setting. Every 10 cycles tubes were briefly spun down in a microcentrifuge to ensure all liquid stayed below the water line in the sonicator. Each sample was then brought to a total volume of 200 μL with RNAse-free water and then precipitated at −20 °C for at least 1 hour or at −80 °C overnight with 600 μL 100% ethanol, 20 μL 3M NaOAc (pH = 5.5), and 2 μL GlycoBlue. RNA was pelleted via centrifugation at 4 °C and 21,000 *g* for 30 min. Pellets were washed twice with 800 mL of ice-cold 70% ethanol, air-dried, and resuspended in 90 μL RNAse-free H_2_O.

ADP-ribose RNA immunoprecipitation was based on a MeRIP-seq protocol for low input samples^69^. 100 μL of Dynabeads Protein G beads were washed 3 times in IP buffer (150 mM NaCl, 10 mM pH 7.5 Tris-HCl, 0.1% NP-40 substitute). 10 μL of Poly/Mono-ADP Ribose rabbit antibody (Cell Signaling Technologies) was added to washed beads resuspended in 500 μL IP buffer and then incubated overnight at 4 °C with end-to-end rotation. Following incubation, antibody conjugated beads were washed twice with IP buffer and then resuspended in 500 μL IP buffer with 20 μg fragmented, rRNA-depleted RNA and 5 μL Superase-In RNAse inhibitor and incubated overnight at 4 °C with end-to-end rotation. Samples were then washed twice with 1 mL IP buffer, twice with 1 mL low salt wash (50 mM NaCl, 10 mM pH 7.5 Tris-HCl, 0.1% NP-40 substitute), and twice with 1 mL high salt wash (500 mM NaCl, 10 mM pH 7.5 Tris-HCl, 0.1% NP-40 substitute). For each wash, beads were incubated in the wash solution for 10 minutes at 4 °C with end-to-end rotation. After the final wash, beads were incubated in 200 μL RLT buffer from the Qiagen RNeasy kit for 2 minutes at room temperature with end-to-end rotation.

Supernatant was separated from the beads using a magnetic rack, transferred to a new tube, and mixed with 200 μL of 100% ethanol. This mixture was passed through a RNeasy MiniElute spin column by centrifugation at 20,000 *g* at 4 °C for 1 minute. Spin columns were then washed once with 500 μL RNeasy RPE buffer and once with 500 μL 80% ethanol with each spin done at 20,000 *g* for 1 minute at 4 °C. Columns were then spun at 20,000 *g* for 5 minutes to remove residual ethanol. RNA was eluted from the column in 15 μL RNAse-free H_2_O with a spin at 20,000 *g* for 5 minutes at 4 °C. RNA yield and integrity was verified using a NanoDrop spectrophotometer and a Novex 6% TBE-urea gel (Invitrogen), respectively.

50-100 ng of pre- and post-IP RNA was then used to make RNA-seq libraries using the NEBNext Ultra II RNA Library Prep Kit for Illumina following the manufacturer’s protocol for use with rRNA Depleted FFPE RNA. Paired-end sequencing of the libraries was performed on a Singular G4 machine at the MIT BioMicroCenter. FASTQ files were then mapped to the MG1655 genome (NC_00913.2), the T4 genome (NC_000866), and the plasmid pKVS45-CmdTAC as previously described^15,70^.

### Library preparation for RNA sequencing

Cells were collected and RNA harvested as described above for infected cells. rRNA was removed using a previously described ribosomal RNA subtraction method^68^. 100 ng of each rRNA depleted RNA sample was then used to make RNA-seq libraries using the NEBNext Ultra II RNA Library Prep Kit for Illumina following the manufacturer’s protocol for use with purified mRNA or rRNA depleted RNA. Paired-end sequencing of the libraries was performed on an Illumina NextSeq 5000 machine at the MIT BioMicroCenter. FASTQ files were then mapped to the MG1655 genome (NC_00913.2), the T4 genome (NC_000866), and the plasmid pKVS45-CmdTAC, as previously described^15,70^.

### Co-immunoprecipitation and LC-MS/MS

Overnight cultures of *+cmdTAC* and *+cmdTA/FLAG-C* cells were back-diluted in 250 mL LB and grown at 37 °C to an OD_600_ of 0.3 and then infected with T4 at an MOI of 10. At 0 and 15 minutes post-infection 64 mL of sample was pelleted by centrifugation at 7,500 *g* for 5 minutes. Pellets were decanted and resuspended in 1 mL of lysis buffer (25 mM Tris-HCL, 150 mM NaCl, 1 mM EDTA, 5% glycerol, 1% Triton X100) supplemented with 1 μL/mL Ready-Lyse Lysozyme (Fischer Scientific), 1 μL/mL benzonase (Sigma), and cOmplete™ Protease Inhibitor Cocktail (Roche) and then flash frozen in liquid nitrogen. Samples were kept in liquid nitrogen until all timepoints were collected. Samples were subjected to two freeze-thaw cycles in liquid nitrogen to ensure complete lysis of cells. Additional lysis buffer was added to samples as needed to normalize sample concentrations by OD_600_. Samples were spun at 20,000 *g* for 10 minutes at 4 °C to pellet any debris. For each sample, 50 μL of Pierce Anti-DYKDDDDK magnetic agarose beads was mixed with 450 μL of lysis buffer and then collected to the side of the tube using a magnetic rack. Beads were then washed twice with 500 μL of lysis buffer. After the final wash, beads were mixed with 1 mL of sample and incubated for 20 minutes at room temperature on an end-to-end rotor. After incubation, beads were washed in wash buffer (1X PBS, 150 mM NaCl) twice and then once with MiliQ H_2_O.

On-bead reduction, trypsin digest, and LC-MS/MS were done as previously described^16^. Detected peptides were mapped to MG1655 and T4 protein sequences and the abundance of proteins were estimated by number of spectrum counts/molecular mass to normalize for protein sizes.

### *In vitro* transcription and translation

In vitro transcription/translation assays were conducted using the PURExpress kit (NEB) according to the manufacturer’s protocol with a 2-hour incubation at 37 °C. Each reaction was supplemented with 1U/µL Riboguard RNase inhibitor (LGC Biosearch Technologies), +/- 1 mM NAD^+^, and protein eluents as indicated. When supplying mRNA as a translation template, primers were used to PCR amplify the DHFR gene from the PURExpress control DHFR plasmid. The PCR amplicon was purified using the DNA Clean & Concentrator Kit (Zymogen). Then, mRNA was synthesized from the PCR template by incubating 300 ng DNA with 200 U T7 RNA polymerase, 0.5 mM NTPs, and 5 mM DTT in a final reaction volume of 40 µL at 37 °C for 4.5 hours. The resulting RNA was purified from the reaction using the RNA Clean and Concentrator Kit (Zymogen) with on-column DNase I treatment. Pure mRNA was then treated with CmdT or control mock purified protein in 1x ADPr buffer (20 mM Tris-HCl pH 8.0 and 150 mM NaCl) with 1 mM NAD^+^ and 1 U/µl Riboguard at 37 °C for 2 hours. RNA was again purified as before, and 1 µg was supplied in the PURExpress reaction for 4 hours at 37 °C. 2.5 µL of the reaction were then denatured in Laemmli buffer and run on a 8-16% polyacrylamide gel by SDS-PAGE and stained with either Brilliant Blue R250 or Coomassie Fluor Orange (Molecular Probes) and visualized on a Bio-Rad ChemiDoc MP imager.

### Protein immunoblotting

Cell cultures were grown overnight and diluted 1:200 in fresh LB containing the appropriate antibiotics. Cultures were grown at 37 °C to mid-exponential phase and then treated with T4 at an MOI of 10, or the appropriate inducers as dictated by the experiment. At various timepoints, cells were pelleted, flash frozen and subsequently resuspended in Laemmli buffer with 2.5% 2-mercaptoethanol in a volume normalized to culture turbidity (100 µl OD600^-1^ ml^-1^). Samples were run by standard SDS-PAGE on 12% polyacrylamide gels. Transfer onto 0.2 µm PVDF membranes was done at 90 V for 40 minutes for CmdA_HA_, and otherwise was done at 100 V for 60 minutes. Membranes were blocked in Tris-buffered saline with 0.05% Tween-20 (TBST) and 5% non-fat milk for 60 minutes at room temperature and incubated with primary monoclonal antibody (1:1000 rabbit anti-FLAG, anti-HA, Cell Signaling Technologies) overnight at 4 °C. Membranes were washed with TBST and incubated with HRP-conjugated goat anti-rabbit IgG (Invitrogen) in blocking buffer for 60 minutes at room temperature. Membranes were again washed and incubated with SuperSignal West Femto Maximum Sensitivity Substrate (Thermo Scientific) before exposure on a Bio-Rad ChemiDoc MP imager. Membranes were stained with Brilliant blue R250 as a loading control.

Non-denaturing blots were performed by lysing cells with a buffer composed of 50% BPER-II (Thermo Scientific), 0.1 mg/mL lysozyme, cOmplete protease inhibitor (Sigma-Aldrich), 6 U DNase-I (NEB) and 3 µL RNase A (NEB) in volumes normalized to OD_600_ of culture samples. Samples were incubated until clear at room temperature and then spun at max speed in a table-top centrifuge for 5 minutes to pellet insoluble material. 6x native loading dye (600 mM Tris-HCl, 50% glycerol, 0.02% bromophenol blue) was added to samples and loaded onto a 12% polyacrylamide Mini-Protean TGX pre-cast gel (Bio-Rad, does not contain SDS). Samples were electrophoresed in 25 mM Tris, 192 mM glycine running buffer at 75 V for 90 minutes. Transfer was conducted onto 0.2 µm PVDF membranes as described in this section at 100V for 1 hour at 4 °C. Blots were processed as described above.

### Burst size determination

Cell cultures of EV and CmdTAC containing strains were grown in LB + 20 µg/mL chloramphenicol in a water bath at 37 °C until OD_600_ measured 0.5. L-tryptophan was then added to 20 µg/mL to each culture to assist adsorption of T4. 100 µL of a 10^7^ PFU/mL T4 stock were added to 9.9 mL of each culture and incubated without shaking for 2 minutes to allow adsorption. Next, for each culture, 100 µL of T4-infected culture from this adsorption flask was added to 9.9 mL LB + 20 µg/mL chloramphenicol (flask A). Flask A was again diluted 1:10 into flask B, and again 1:10 into flask C. 500 µl from flask A was added to 200 µL ice-cold chloroform and vortexed for 10 seconds. Viable PFUs from this chloroform-treated sample represent unadsorbed phage (adsorption control). Next, 100 µL from each flask A (time 0 sample) or the adsorption control was mixed with 3.5 mL LB 0.5% agar maintained at 50 °C to which was added 50 µL of an overnight culture of indicator strain. This mixture was vortexed briefly and overlayed onto LB + 20 µg/mL chloramphenicol + 1.2% agar plates. All flasks were then left to incubate in a shaking water bath at 30 °C. After 60 minutes, 100 µL from flask C of the empty vector strain and flask A of the +*cmdTAC* strain were overlayed with indicator strain on agar plates. After overnight incubation at 37 °C, plaques were enumerated, and normalized to the adsorption control. Burst size was recorded as the number of plaques from each plate multiplied by their dilution factor, and then divided by the number of plaques at time 0.

### T4 genome engineering and evolution

The evolution of T4 on *cmdTAC* containing cells was performed as described previously^71^ for 4 rounds, resulting in the *alt.-3* C-terminal extensions in all five replicates. To generate *alt.-3* mutants for further evolution experiments, T4 stock was overlayed onto strains containing Cas9 and spacers directed toward *alt.-3* or a control plasmid with no spacer (ML4233 and ML4234). The number of plaques formed on the spacer containing strain was compared to the control to determine if there was any selection imposed by the spacer. Despite attempting 8 potential spacers, no selection was observed. To mitigate this, we repeated the experiment with T4 Δ*agt* Δ*bgt* (DNA contains non-glucosylated, 5-hydroxymethyl cytosine) on an *E. coli* Δ*mcrA* Δ*mcrBC* background required for viability of T4 Δ*agt* Δ*bgt*. This T4 formed fewer plaques in the presence of the *alt.-3* spacer, suggesting that selection for *alt.-3* mutants was imposed in this condition. The *alt.-3* region was PCR amplified and Sanger sequenced from plaques that were able to form on the spacer containing strain. Of those plaques, we isolated a strain that harbored a mutation encompassing nearly the entire open reading frame of *alt.-3*. The T4 Δ*alt.-3* strain was propagated in the presence of the spacer and stored as a stock at 4 °C. Evolution of this T4 strain on *cmdTAC-*containing cells was conducted the same as before, for 17 rounds, without observing mutations that increased plaquing ability.

### Radiolabel incorporation assays

Overnight cultures of +*cmdTAC* and +*cmdT*AC* cells were back-diluted in LB + 20 µg/mL chloramphenicol and grown at 37 °C to an OD_600_ between 0.2 and 0.3. An aliquot of each culture was collected before T4 infection at an MOI of 10 and again at each indicated timepoint (t=10, 20, 30, 40 minutes post-infection). Each collected sample was incubated with EasyTag EXPRESS-^35^S protein labelling mix (Perkin Elmer) at 23 µCi mL^-l^ for 2 min at 37 °C. Labelling was chased with an unlabeled cysteine/methionine mixture at 5 mM and then samples precipitated in 13% w/v ice-cold TCA. Samples were pelleted by centrifugation at 16,000 *g* for 10 minutes at 4 °C, washed twice with 500 µL ice-cold acetone, and then resuspended in resuspension buffer (100mM Tris pH 11.0, 3% w/v SDS). Samples were run on 4-20% SDS-PAGE gels (BioRad), the gels incubated in Gel-Dry Drying Solution (Invitrogen) for 10 minutes, and then dried on a vacuum gel dryer for 2 hr at 80 °C. Dried gels were exposed to a phosphorimaging screen overnight before imaging on an Amersham Typhoon imager.

### Protein purification

5 mL of cultures of ML4207 and ML4232 were grown overnight at 37 °C in LB + 0.2% glucose. The following day, 5 mL of each culture was washed of glucose twice and used to inoculate 495 mL of LB + 25 µg/mL kanamycin. After one hour of additional growth, arabinose was added to a final concentration of 0.2%. Cultures were grown an additional 95 minutes, pelleted, washed with H_2_O, again pelleted, and flash frozen in liquid N_2_. The following day, cell pellets were resuspended in 4 mL lysis buffer (50 mM Tris pH 7.5, 500 mM NaCl, 0.05% Tween-20, EDTA-free protease inhibitor, 0.5 mM PMSF, 0.5 mg/mL lysozyme, 5 mM imidazole, and 5% glycerol) on ice. Cells were then lysed by sonication in a Bioruptor 300 for two rounds of 10 cycles each, high setting, 30 s on/ 30 s off. 1 mL of Ni-NTA agarose resin (Qiagen, 0.5 mL bed volume) was equilibrated in lysis buffer. Cell lysate was clarified by centrifugation then incubated with the Ni resin for one hour at 4 °C with gentle rocking. The following steps were conducted at 4 °C. The resin was then passed through a 10 mL chromatography column and then washed 5x with 2.5 mL of wash buffer (same as lysis buffer but without lysozyme, and imidazole at 20 mM). Protein was then eluted 5x with 2.5 mL elution buffer (wash buffer with imidazole concentration at 300 mM). Eluted proteins were buffer-exchanged into Tris pH 7.4 using Micro Bio-Spin chromatography columns (Bio-Rad) and concentrated using Amicon Ultra 0.5 mL centrifugal filters with a 3 kD cutoff.

### *In vitro* ADP-ribosylation by CmdT

A typical reaction was assembled on ice as follows. To a buffer composed of 20 mM Tris-HCl pH 8.0 and 150 mM NaCl, we added 1U/µL Riboguard RNase inhibitor, 1 mM NAD^+^ (NEB), 4 µg of DNA or RNA oligo, and protein at the concentrations indicated. The reactions were then incubated in a thermocycler at 37 °C. To stop the reaction, an equal volume of 2x 6M urea sample buffer (Novex) was added. RNA was denatured at 95 °C for 10 minutes and then immediately placed on ice. 1 µg of RNA samples were then subject to electrophoresis in 15% polyacrylamide TBE-urea gels at 180 V for 75 minutes. Gels were stained both with SYBR-Gold and with a concentrated solution of 0.2% methylene blue in 0.1x TBE buffer for 15 minutes, de-stained with several changes of H_2_O and imaged.

### *In vitro* ADP-ribose RNA pulldown

20 µg of total RNA was ADP-ribosylated with CmdT as described above with 0.25 mM 6-Biotin-17-NAD^+^ for 4 hours at 37 °C and then continued at 4 °C overnight. Two control reactions were set up identically except with mock purified protein, or with standard NAD^+^ in place of 6-Biotin-17-NAD^+^. 10 µg of each reaction was kept at −80 °C as the pre-pulldown sample. The remaining 10 µg were incubated with streptavidin conjugated superparamagnetic beads (Dynabeads MyOne Streptavidin C1) following the manufacturer’s protocol. RNA was stripped from the beads by addition of 0.5 mL Trizol and incubation at 25 °C for 15 minutes on a thermomixer at 1000 rpm. Beads were then precipitated with a magnet and 100 µL of chloroform were added. The phases were separated by centrifugation at 14,000 *g* for 15 minutes. Finally, the aqueous phase was purified using the RNA Clean and Concentrate Kit (Zymogen). Pre- and post-pulldown samples were electrophoresed on a 6% TBE-urea gel for 45 minutes, stained with SYBR-gold and imaged. The samples from pre- and post-pulldown reactions containing 6-Biotin-17-NAD+ and CmdT were subject to RNA sequencing as described in this section, but without rRNA-depletion.

### HPLC analysis of ribonucleosides

10 µg each of no-U and no-C RNA oligos (Figure 6D) were subjected to ADP-ribosylation as described above. Controls were included in which purified CmdT was replaced by a mock purification, or in which NAD^+^ was omitted. Next, samples were split and treated with either 100 U Nuclease P1 (NEB) and 10 U Antarctic phosphatase, or, the same with the addition of 1 U Phosphodiesterase I from Crotalus adamanteus venom (SVPD, Millipore Sigma). Reactions were incubated in digest buffer (25 mM Tris-HCl pH 7.6, 50 mM NaCl_2_, 1 mM ZnCl_2_, and 10 mM MgCl_2_) at 37 °C for 3.5 hours in a total volume of 110 µL. 100 µL of digested and dephosphorylated nucleosides (10 µg) were injected onto a Vydac C18 4.6 x 250 mm reverse phase silica column (218TP54) equilibrated with equilibrated with 90% buffer A (0.1M triethylammonium acetate (TEAA), pH 7.0)/10% buffer B (0.1M TEAA, 20% acetonitrile, pH 7.0). HPLC was run with a mobile phase gradient composed of buffer A and B, 10-60% B from 1-21 minutes and 60-97% B from 21-26 minutes at a flow rate of 1 mL/minute. Analytes were measured at A_254_. On a replicate run, samples without SVPD treatment were collected as fractions and relevant fractions were lyophilized. The samples were then resuspended in digest buffer, and again incubated for 3.5 hours with 10 U Antarctic phosphatase and 1 U SVPD and analyzed by HPLC as described above.

### Mass spectrometry of modified nucleosides

Fractions collected from HPLC analysis were dried in a speed-vac and resuspended in 200 µL of 50% acetonitrile in 0.1% formic acid. The fractions, or a buffer blank, were directly infused by syringe pump into a Thermo Q Exactive with an API source and electrospray ionization probe at a flow rate of 5 uL/minute. The instrument was operated in positive ion mode. MS/MS was conducted at collision energies of both 25 and 40 CE. Instrument parameters were as follows: spray voltage, 3.8 kV; capillary tube temperature, 280 °C; sheath gas, 20; auxiliary gas, 5; sweep gas, 5.

### Bioinformatic analyses

The CmdT sequence logo was generated from the CmdT hmm file from DefenseFinder^66^ using skylign.org.

HMM profiles of CmdT and CmdC were downloaded from DefenseFinder^66^ and searched against the RefSeq non-redundant protein database using hmmscan and default parameters. Protein hits were then identified in all available RefSeq bacterial genomes and CmdTAC was called if both CmdT and CmdC were present within two proteins of each other in the genome. CmdA was not included in calling as it both has higher sequence variability and is often unannotated in clearly homologous systems. All datasets were downloaded in July 2023. The complete taxonomic lineage of refseq genomes was created and filtered to include bacteria of current interest (genera with > 1000 sequenced genomes). A taxonomic relationship of these genera was produced with NCBI Common Tree, and presence/absence was recorded from the taxonomic profiles of the CmdT/C hmmscan.

### Structural Predictions

Predictions of the structures of individual components and CmdTAC as a complex were done using AlphaFold2 (ref^25^) with the multimer module and default parameters on the reduced database with 1 prediction generated per model. Structural homology searches based on the AlphaFold2 predicted structures of CmdT and CmdC were done using Foldseek^64^.

### RNA-sequencing and RNA-IP sequencing read mapping

FASTQ files for each sample were trimmed using cutadapt^57^ (version 1.15) and then mapped to the *E. coli* MG1655 genome (NC_00913.2), the T4 genome (NC_000866), and the plasmid pKVS45-CmdTAC using bowtie2 (ref^56^) (version 2.3.4.1) with the following arguments: −D 20, −I 40, −X 300, −R 3, −N 0, −L 20, −i S,1,0.50. Sam files generated from bowtie2 mapping were then converted to bam files using samtools^58^ (version 1.7) and then further converted to numpy arrays using the genomearray3 python library^65^ for use in downstream analyses. For *in vivo* RIP-seq analyses only highly expressed transcripts as determined by transcripts with an RNA TPM for both replicates greater than or equal to the minimum mean TPM of any T4 transcript were used for the generation of figure 5D. For statistical comparisons of TPM ratios between RNA types a Welch’s t-test was used.

### Co-IP LC-MS/MS Analysis

Spectral counts were calculated for each protein in the *E. coli* MG1655 and T4 proteomes and the CmdTAC system and high confidence hits assessed in downstream analyses. The ratio of spectral counts between the FLAG-CmdC pulldown and untagged CmdC pulldown at each timepoint were used to generate Fig. 3 with a pseudocount added to each count.

---

**Figure S1.**
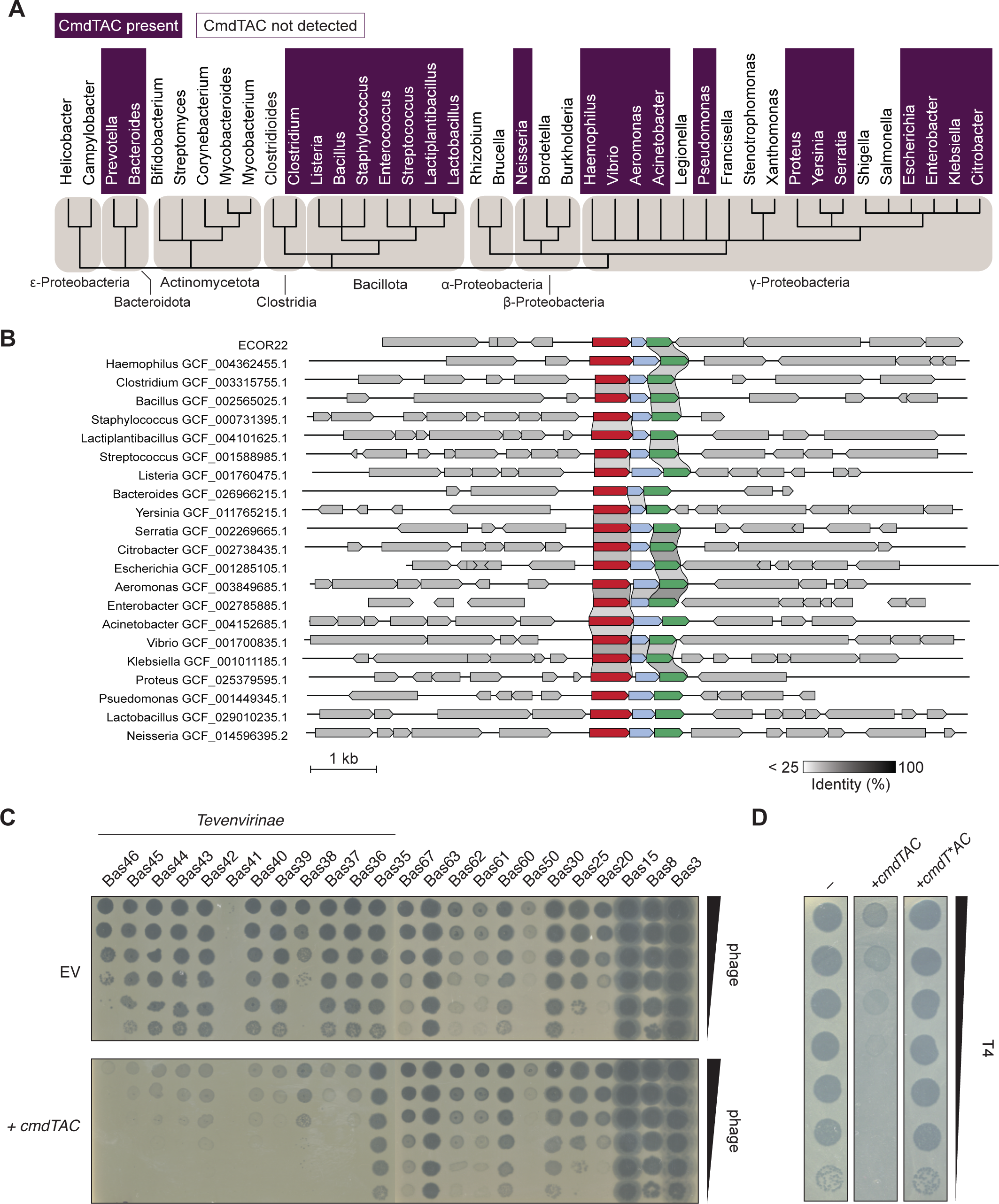
Taxonomic distribution of *cmdTAC* and efficiency of plaquing for BASEL and T-even phages. (A) Presence or absence of CmdTAC homologs in bacterial genera with > 1000 sequenced genomes (see Methods). (B) Examples of CmdTAC homologs found in diverse bacterial species. Grey bars between genes capture percent identity, defined by the color bar below. (C) Plaquing of the phages indicated on *E. coli* K12 harboring *cmdTAC* or an empty vector control. Data used to generate EOP data in Fig. 1C. (D) Plaquing of T4 phage on EV, *cmdTAC*, or *cmdT*AC*.

**Figure S2.**
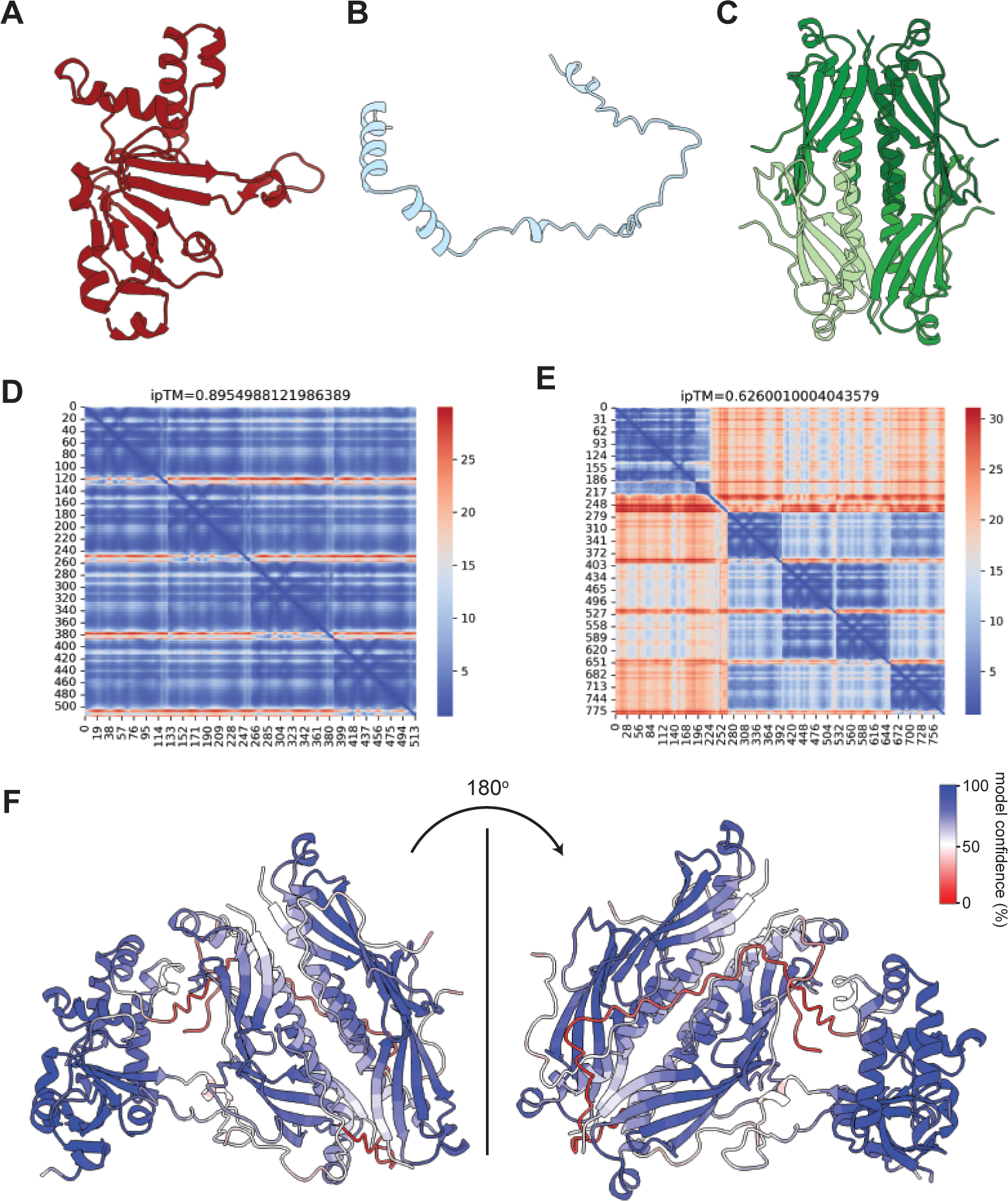
AlphaFold2-based prediction of CmdTAC structures. (A-C) AlphaFold2 predicted structures of CmdT (A), CmdA (B), and tetrameric CmdC (C). (D) PAE plot and per-residue model confidence score (pLDDT) for AlphaFold2-predicted CmdC tetramer. (E) PAE plot and per-residue model confidence score (pLDDT) for the AlphaFold2-predicted complex of CmdTAC. (F) Ribbon diagrams of predicted CmdTAC complex, color-coded based on per-residue model confidence score (pLDDT).

**Figure S3.**
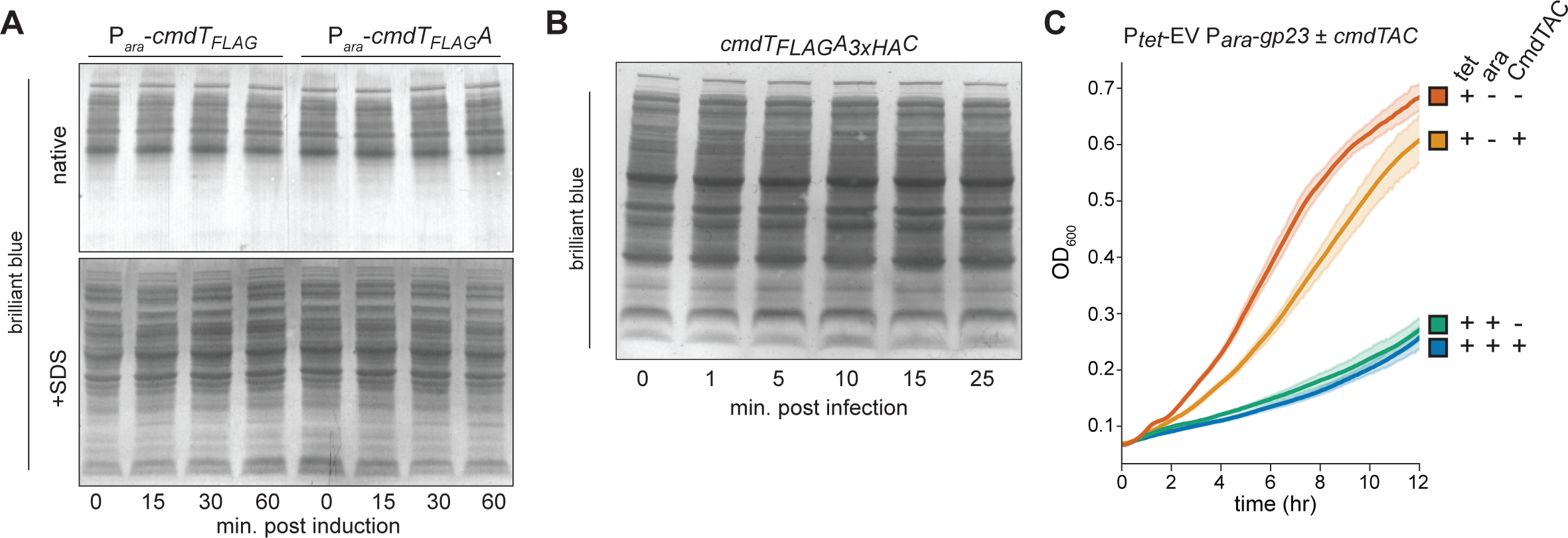
Loading controls of for immunoblots, and effects of producing Gp23 alone on cell growth. (A-B) Coomassie stained gels used as loading controls for immunoblots shown in Fig. 2B and 2E, respectively. (C) Growth curves for uninfected *E. coli* cells producing CmdTAC from its native promoter and induced to produce Gp23 (T4 major capsid protein), as indicated, all without Gp31, the T4 chaperonin for Gp23. Data are the average of 3 biological replicates.

**Figure S4.**
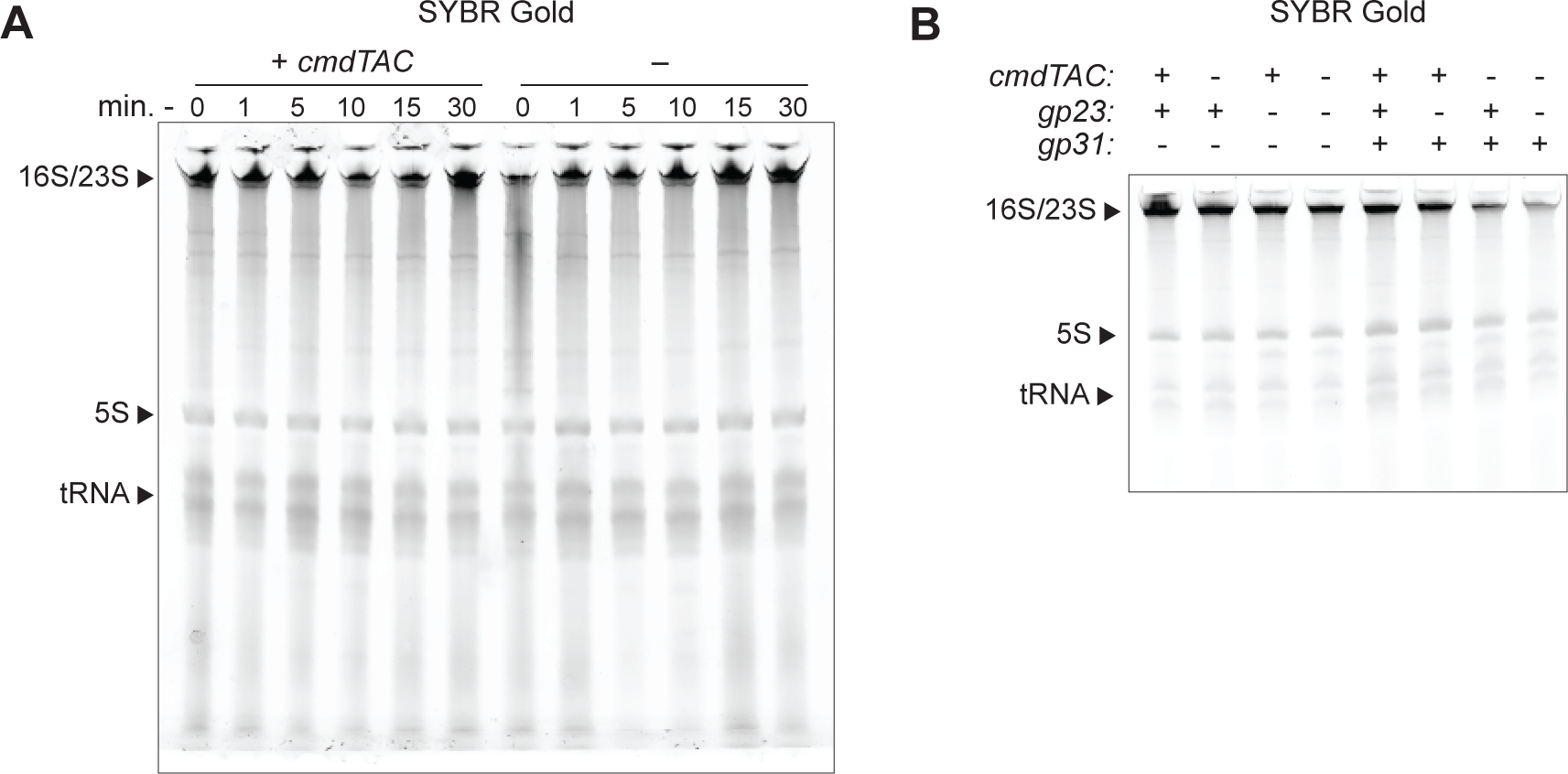
Total RNA staining of TBE-Urea gels used in immuno-northern blotting experiments. (A-B) Total RNA run on TBE-Urea gel stained with SYBR gold used in immune-northern blotting during T4 infection of cells harboring *cmdTAC* (A) or during ectopic expression of Gp23 and Gp31 in uninfected cells harboring *cmdTAC* (B) as shown in Figures 4D and 4E, respectively.

**Figure S5.**
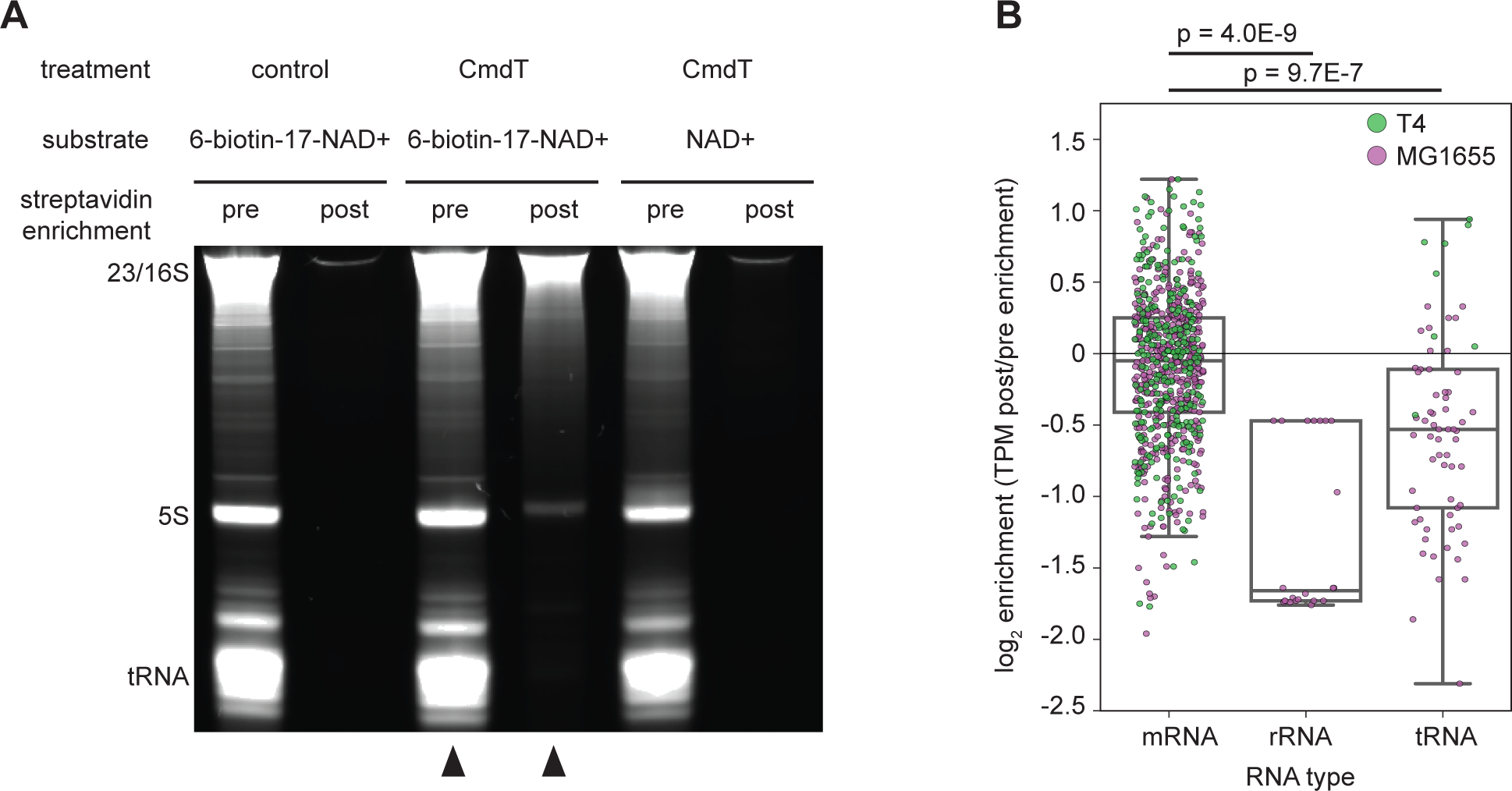
In vitro ADP-ribosylation and RNA-precipitation sequencing with 6-biotin-17-NAD^+^. (A) TBE-urea gel showing total RNA from T4-infected cells after treatment +/- CmdT and with 6-biotin-17-NAD^+^ or NAD^+^. RNA is shown pre- and post-streptavidin enrichment, with arrows indicating samples that were sequenced. (B) Ratio of TPM after streptavidin enrichment of biotinylated RNA to TPM pre-enrichment for each transcript. Transcripts are separated by RNA type and T4 vs MG1655 origin. Note that the T4 genome does not contain rRNA. Welch’s t-test p-values are indicated.

**Figure S6.**
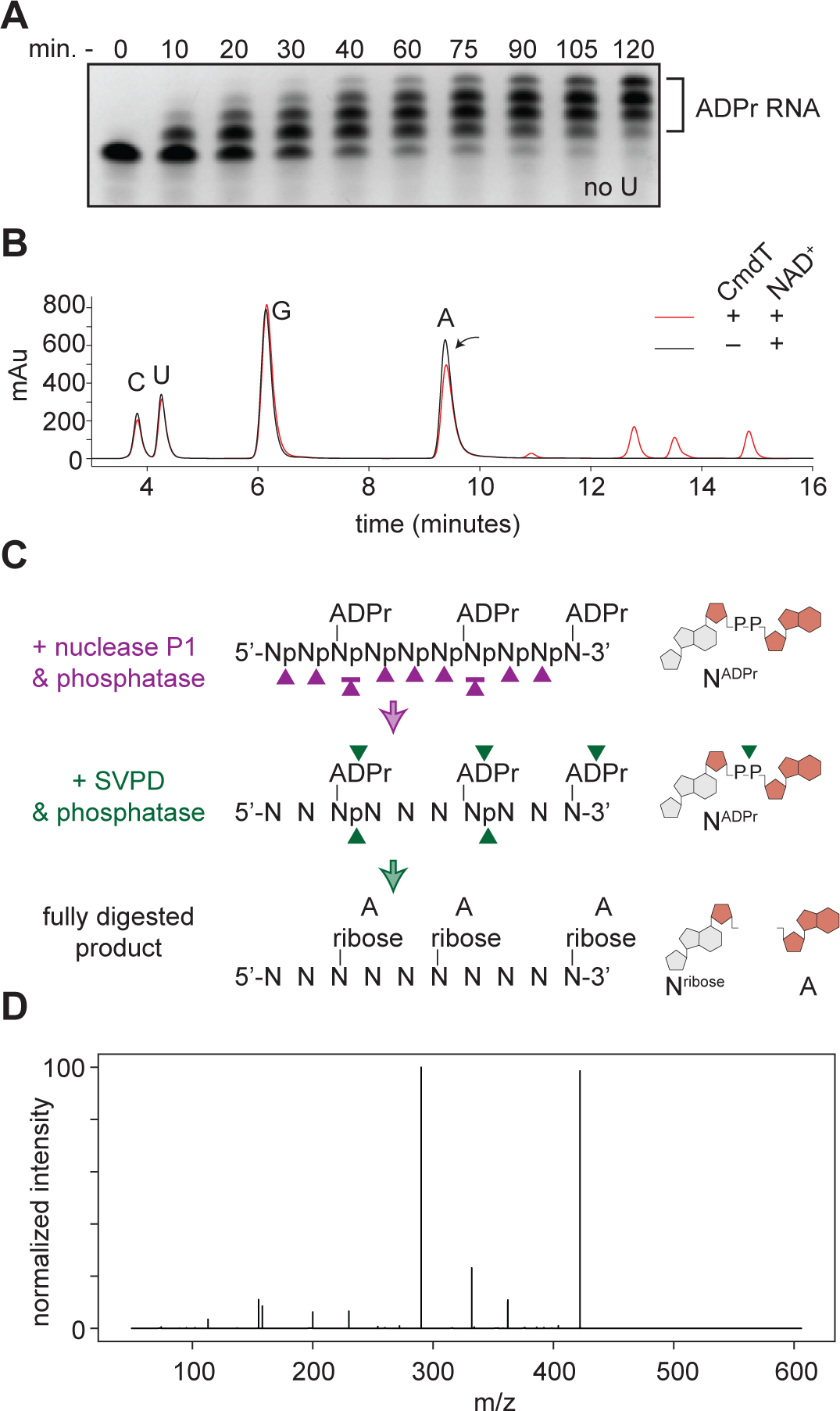
Additional analyses of ADP-ribosylation of RNA by CmdT. (A) The no-U oligo (see Fig. 6D) incubated with CmdT for the times indicated, with all reactions visualized as in Fig. 6. (B) Overlay of HPLC traces from Fig. 6F derived from analysis of nucleosides isolated following incubation of the no-U and no-C RNA oligos with CmdT and NAD^+^ (or without NAD^+^, as indicated) and then treated with nuclease P1 and antarctic phosphatase. Arrow highlights the decrease in adenosine for reaction containing CmdT and NAD^+^. Peaks corresponding to A, C, G, and U nucleosides are marked. (C) Schematic summarizing enzymatic digestion activities of nuclease P1, antarctic phosphatase, and snake venom phosphodiesterase on ADP-ribosylated RNA oligos. (D) MS/MS fragmentation of the A-ribose sodium adduct. Collision energy = 20 eV, as used previously^11^. Note the absence of a ribose-ribose peak at m/z = 287.

**Figure S7.**
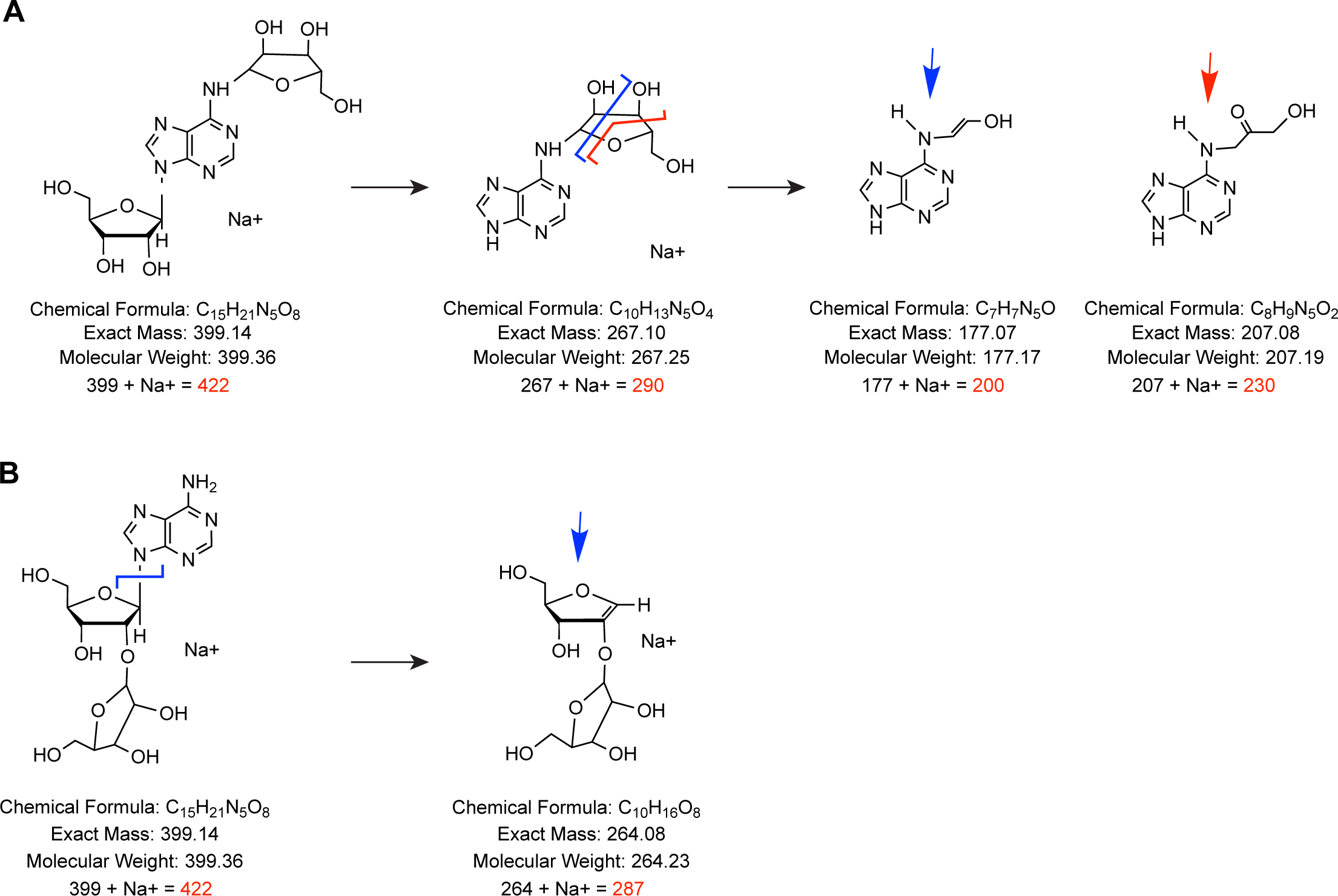
Mass spectrometry analysis of CmdT-dependent ADP-ribosylated adenosine. (A) Predicted fragmentation of adenosine ribosylated on the N6 position (far left) to produce ribosylated adenine (second from left), which is predicted to further fragment into two species (right), with m/z values of 200 and 230, corresponding to the peaks seen in Fig. 6J. (B) Predicted fragmentation of adenosine ribosylated on the 2’-OH (left) to produce the ribosylated adenine (right), with an m/z value of 287. This peak was not seen in Fig. 6J, but was seen with prior analysis of RhsP2 (ref^11^).

## Notes

### Competing Interest Statement

The authors have declared no competing interest.

